# Exercise training improves sarcopenic muscle function via restoration of mitochondrial quality control

**DOI:** 10.1101/2025.11.03.686168

**Authors:** Isabelle Alldritt, James Sligar, Dean Campelj, Camila S. Padilha, Elya Ritenis, Theo Mantamadiotis, Samuel Widodo, Marija Dinevska, Joachim Nielsen, Ashleigh M. Philp, Andrew Philp

## Abstract

Mitophagy is an essential component of the mitochondrial quality control program, maintaining mitochondrial homeostasis in metabolic tissues such as skeletal muscle. With age, it is thought that mitochondrial quality control becomes dysregulated, leading to the progression of age-associated diseases such as sarcopenia. Exercise is known to enhance skeletal muscle mitochondrial health and may be an effective intervention to prevent sarcopenia, however the role of mitophagy in this process is unknown. Utilising mitophagy reporter mice (mito-QC), we assessed adaptations in skeletal muscle mitophagy in response to increased age (3-26 months) and following an 8-week endurance exercise training period. Immunofluorescent imaging revealed that ageing led to an accumulation of mitolysosomes in sarcopenic old mice indicative of increased mitophagy, an adaptive response that was reversed by exercise training. In parallel to reducing age-associated mitophagy, exercise training increased mitochondrial respiratory capacity and improved muscle strength, suggesting that alterations in mitochondrial quality control led to improvements in skeletal muscle function. Exercise-mediated alterations in mitophagy were accompanied by increases in BNIP3, FUNDC1 and BCL2L13 protein content post training. Collectively our data suggests that sarcopenia leads to dysregulation of mitophagy in skeletal muscle. Restoring mitophagy balance with exercise training leads to improvements in mitochondrial respiration and skeletal muscle strength, identifying a novel cellular mechanism to explain the benefits of exercise training in old age.

## Introduction

Increased age is associated with a change in mitochondrial morphology^1^, loss of mitochondrial function, and decreased total mitochondrial content^2^. These processes are regulated at the cellular level by the mitochondrial quality control program^3^, which orchestrates mitochondrial homeostasis via fluctuations in biogenesis, dynamics and mitophagy. As skeletal muscle has a high energetic demand, it is particularly susceptible to alterations in mitochondrial quality control^4^, with alterations in this program proposed to play a key role in the aetiology of sarcopenia, the loss of skeletal muscle mass and function commonly associated with age^5^.

While age-related declines in mitochondrial biogenesis are well reported^6–8^, mitophagy is poorly understood in the context of ageing and there is disagreement as to whether mitophagy increases^9–11^ or declines^12–14^ in aged tissue. Such discrepancies may be attributed to the timing of collection and the use of static markers which measure aspects of the mitophagy machinery rather than mitophagy itself. The use of global and conditional mitophagy reporter mouse models^15–17^ circumvents this technical limitation, by allowing for the direct quantification of mitophagy in mammalian systems. Using these models, increased mitophagy has been linked to exercise adaptation^18^ and tissue ageing^19^ and muscle mitochondrial pathologies^20^.

It is well established that exercise has a potent effect on mitochondrial quality and function^21^, even in older sedentary individuals^22^. This may be mediated through the induction of mitophagy in response to the exercise stimulus^23,24^. Here, we use the mito-QC mitophagy reporter mouse model to quantify changes in skeletal muscle mitophagy over a 26-month period in male and female mice and following exercise training in old mice. Our results demonstrate that ageing and the progression of sarcopenia leads to an increase in skeletal muscle mitophagy which alters the balance mitochondrial quality control. Exercise training in old mice leads to a restoration of mitophagy to young levels, in parallel to increases in mitochondrial function and muscle strength. Collectively our data supports a role for mitophagy in sarcopenia progression and highlights the therapeutic value of targeting this pathway in the treatment and prevention of sarcopenia.

## Methods

### Experimental model and subject details

#### Mice

All experiments were carried out with the approval of the Garvan Institute of Medical Research/St Vincent’s Hospital Animal Ethics Committee (application number 21_20) following the requirements set by the state of New South Wales and the National Health and Medical Research Council of Australia. All animal experiments were conducted by trained personnel and were performed in accordance with the Australian code of practice for the care and use of animals for scientific experimentation. Sample size was determined based on our previous work^25^ utilising the mean response from pre (PRE: 1.001 ± 0.12 vs POST: 2.73 ± 0.48; Mean ± SEM) for COX-V (representing the smallest response in COX subunit protein content) following exercise training. Based on this response, a sample size of 8 was required in each group to detect a training induced increase in mitochondrial protein content (α = .05, 1-β = .90, effect size f = .18). Factoring in variance in training adaptation, we chose to study 9 mice per group/time-point.

Mito-QC mice^15^ were provided by Professor Ian Ganley (School of Life Sciences, The University of Dundee, Dundee, Scotland). Briefly, a CAG promoter cassette and the open reading frame for the mCherry-GFP-FIS1 fusion protein including a Kozak sequence (GCCACC) were inserted into the mouse *Rosa26* locus, producing mitochondria which fluoresce both green and red. The pH shift induced by the formation of a mitolysosome during mitophagy quenches GFP signal but does not affect mCherry expression. Puncta positive for mCherry expression but negative for GFP expression can be defined as mitolysosomes. Mice were bred and maintained on a C57BL/6 background at the Australian Bioresource facility, Moss Vale, NSW. All mice were housed for at least one week prior to initiation of experimentation. Mice were grouped housed in a temperature-controlled environment (22°C) with a 12-hour light/dark cycle. Male and female mito-QC mice were aged to 3, 12, 24 or 26 months (n=9/group/sex). At 24 months, mice were randomly allocated to either sedentary or exercised training groups.

### Method details

#### Body composition analysis

Body weight, lean mass and fat mass were obtained the week prior to tissue collection at each experimental timepoint using an EchoMRI (EchoMRI LLC, Houston, USA) as previously described^26^.

#### Exercise training intervention

Male and female (n=9/sex) 24-month-old mice underwent a custom 8-week progressive treadmill intervention on a custom-built treadmill (Columbus instruments, USA), completing three sessions of 60 minutes per week (Extended Data Figure 1). Each session consisted of 5 minutes of acclimatisation, 30 minutes of low intensity exercise, and 4 rotations between 5-minute intervals at alternating high and low intensity, followed by a 5-minute cooldown period. Exercise training began at 8.5 m/min (week 1) and was increased by 1 m/min each week.

#### Tissue collection

One hour prior to tissue collection, food was removed, and body weights recorded. Mice were anesthetized with 2% isoflurane and tissues harvested and weighed. Terminal blood samples were taken via cardiac puncture followed immediately by euthanasia via cervical dislocation. Tissues were either snap frozen in liquid nitrogen and stored at -80°C until use or immediately placed in BIOPS buffer for use in high-resolution respirometry.

#### High resolution respirometry

Before measurement of respirometry, muscle fibres were removed of connective tissue and separated into bundles using forceps. Fibre bundles were placed into 2ml BIOPS buffer with 50µg/µl saponin to permeabilise fibres. Samples were gently rocked at 4°C for 30 minutes, then placed into 2ml of MiR05 buffer and gently rocked for 10 minutes at 4°C. To prevent further permeabilization, bundles were placed in fresh MiR05 buffer. Mitochondrial respiration was measured using high-resolution respirometry Oxygraph-2k (Oroboros Instruments, Austria) as previously described^26^. Chambers were maintained at a constant temperature of 37°C and oxygen concentration was maintained between 150 and 220 µM. Bundles totalling 2.5-5mg muscle were blotted dry and added to each chamber. Pyruvate (10mM) and malate (2mM) were added to assess complex I-related leak (CI_L_). ADP was titrated in stepwise increments (0.1-6M) followed by addition of glutamate (10mM) to assess phosphorylating respiration (CI_P_). Succinate (10mM) was added to assess respiration support via complex II (CI + II_P_). Cytochrome c (10µM) was added to test outer mitochondrial membrane integrity, with the threshold for acceptance set at within 10% of succinate. Carbonyl cyanide 3-chlorophenylhydrazone (CCCP) was titrated in a stepwise manner to reach a final concentration of 0.5µM. Finally, antimycin A (2.5µM) was injected to determine non-mitochondrial oxygen consumption.

#### Mitophagy immunofluorescence quantification

Immunofluorescent images were acquired using a Leica SP8 (Leica Microsystems, Germany) confocal microscope. For longitudinal imaging of tissue, images were acquired at 100x magnification. A minimum of 3 whole fibres were captured per image. For cross-sectional imaging of tissue, images were acquired at 40x magnification with 20-35 whole fibres captured per image. A composite of image from green and red channels was generated for analysis. Mitophagy was quantified using the *mito-QC Counter* macro for FIJI^27^ which examines the ratio of mCherry to GFP intensity of individual pixels within a defined region of interest. Sites with a high mCherry:GFP ratio were defined as sites of mitolysosome formation.

To optimise the *mito-QC Counter* macro for use with skeletal muscle tissue, the ratio threshold, smoothing radius, and standard deviation above red mean parameters were adjusted until quantification of mitophagy was accurate when compared to a count of mitolysosomes determined from manual selection. Following optimisation, ratio threshold value was set at 0.8, smoothing radius was set at 1, and standard deviation above red mean was set at 10,000 (Supplementary Methods).

#### Selection of mitochondrial aggregates

Structures that fluoresced green in the composite image, were wider than the size of a mitochondria from visual inspection of the image, had clear boundaries to the fibre, and obstructed the view of the mitochondrial network in the fibre were identified as mitochondrial aggregates and size was measured manually. To exclude outliers, aggregates were removed from further analysis if size was greater than two standard deviations above the mean.

#### Transmission electron microscopy

A small specimen of muscle was fixed in 2.5% glutaraldehyde in 0.1M sodium cacodylate buffer for 24 hours at 5°C and washed in sodium cacodylate buffer. Fixed samples were processed as described elsewhere^28^. Briefly, samples were post-fixed with 1% osmium tetroxide and 1.5% potassium ferrocyanide in 0.1M sodium cacodylate buffer, rinsed in the buffer, dehydrated by a graded series of alcohol, infiltrated with graded mixtures of propylene oxide and Epon, and embedded in 100% fresh Epon and polymerised at 60°C for 48 hours. Two 60nm sections were cut by an ultramicrotome (Leica Microsystems, Germany) interspaced by 150µm in depth of the Epon block and contrasted by uranyl acetate and lead citrate. Prior to imaging, samples were anonymised. Sections were imaged in a pre-calibrated EM 208 transmission electron microscope (Philips, The Netherlands) using a Megaview III FW camera (Olympus Soft Imaging Solutions, Germany). Each sample was photographed by a systemic but random protocol that ensured unbiased results. Equal number of images of subsarcolemmal (SS) and intermyofibrillar (IMF) regions of each fibre were acquired. Images were analysed by manually outlining mitochondria present in each fibre and measuring length of fibre using the “interpolated polygon” and “polyline” functions in RADIUS software (EMSIS GmbH, Germany).

#### Ex-vivo muscle excitation contraction coupling

To assess muscle contractile function, excised extensor digitorum longus (EDL) and soleus (SOL) muscles were placed into individual organ baths (Danish Myogenic Technology, Denmark) filled with a modified Krebs-Henseleit Ringer’s solution as previously described^29^. Each bath was supplied with carbogen (5% CO_2_/95% O_2_) and maintained at a constant temperature of 30°C and pH of 7.4. The proximal end of the muscle was attached to a force transducer which had been previously calibrated, and the distal end was fixed to a micromanipulator with stimulating electrodes. Absolute twitch and tetanic force production were established, followed by a force frequency relationship as described elsewhere^30^. Force production per muscle cross-sectional area (specific force production) was calculated according to Brooks and Faulkner^31^. To investigate muscle fatiguability, the EDL was tetanically stimulated for 350ms at 100Hz every 4 seconds and the SOL was tetanically stimulated for 500ms at 80Hz every 2 seconds for a total duration of 3 minutes. Data were collected and analysed using LabChart Pro Version 8.0 (AD Instruments, NZ).

#### Western blotting

Approximately 20mg muscle tissue was powdered and homogenised in sucrose lysis buffer with protease inhibitor tablet as described previously^26^. Samples were centrifuged at 18,000 xg for 20 minutes at 4°C. Protein concentration in the supernatant was determined by DC protein assay. Supernatant was added to 25µl 4x Laemmli buffer with B-mercaptoethanol as a reducing agent and lysis buffer such that final protein concentration was 40µg/µl. Samples were denatured by boiling at 95°C for 5 minutes. 20µg protein was loaded onto 10% SDS-PAGE gel and transferred onto a nitrocellulose membrane. To confirm protein transfer, membranes were incubated in Ponceau S dye solution (0.1% (w/v) in 5% acetic acid). Blots were blocked for 1 hour with 3% milk in TBST at room temperature and incubated overnight at 4°C with the primary antibody (1:1000 dilution in TBST). Membranes were washed with TBST and incubated with anti-rabbit IgG secondary antibody conjugated with horseradish peroxidase (HRP, 1:10,000 dilution in TBST) for 1 hour at room temperature. Proteins were detected using enhanced chemiluminescence (ECL) HRP detection reagents. Images were captured using the ChemiDoc TOUCH Imager (Bio-Rad, USA) and protein content was quantified using ImageJ (NIH, USA).

#### Antibodies

All primary antibodies were used at a 1:1000 dilution in buffers following manufacturer instructions. Parkin (2132S) was from New England Biolabs. FUNDC1 (49250), BNIP3 (44060), BCL2L13 (61974), Optineurin (58981), FKBP8 (18582), NIX (12396) and STING (50494) were acquired from Cell Signalling Technology. Anti-rabbit secondary antibody (#7404) acquired from Cell Signalling Technology was used at a 1:10,000 dilution in TBS-T.

#### Metabolomics sample preparation

Metabolites were extracted from skeletal muscle tissue by a dual phase methanol-chloroform-water method^32^. Approximately 20mg tissue was chipped and powdered. 1.6µl MeOH/mg tissue and 0.53µl ddH_2_O/mg tissue were added for homogenisation. 0.8µl CHCl_3_/mg tissue was added and samples were centrifuged 4°C for 5 minutes at 2500 xg. 0.8µl CHCl_3_/mg tissue and 0.91µl ddH_2_O/mg tissue were added to supernatant. Samples were agitated for 2 minutes before being incubated at 4°C for 10 minutes, then centrifuged at 17,000 xg for 10 minutes at 4°C. Upper and lower phases were removed to separate Eppendorf tubes and dried under a vacuum. Prior to analysis, samples were suspended in 75:25 MeCN:ddH2O and 75:25 IPA:ddH2O for HILIC and reverse phase chromatography, respectively. 10 µl was removed from each suspended plasma sample to a clean Eppendorf tube to create a representative quality control (QC) sample.

#### Liquid chromatography-mass spectrometry analysis of metabolites

Metabolomics analysis was performed on an Ultimate 3000 UHPLC pump and autosampler coupled to a Q-Exactive mass spectrometer (Thermo Fisher Scientific, USA) operated in positive and negative ionisation modes. For HILIC chromatography, an InfinityLab Poroshell 120 HILIC-Z column (Agilent) was used with mobile phase A 10mM ammonium formate in 95% acetonitrile with 0.1% formic acid and mobile phase B 10mM ammonium formate in 50% acetonitrile with 0.1% formic acid. For reverse phase chromatography, a Zorbax SB-Aq RRHD column (Agilent) was used with mobile phase A ddH2O with 0.1% formic acid and mobile phase B methanol with 0.1% formic acid. 5µl sample was injected. Injection order was randomised to avoid bias. 8 pooled QC samples were injected at the start of the run for column conditioning, followed by an injection every 10 biological samples.

Data was converted to an open mzXML format using MS Convert (ProteoWizard, USA) and processed using inhouse R scripts^33^ utilising the XCMS package^34^. Chromatograms were extracted using the centWave algorithm. Tolerance was set at 20ppm. Retention time alignment was performed using the Obiwarp method with a bin size of 0.6. Bandwidth parameters for correspondence analysis were determined manually for each ion mode and polarity. Following correspondence, missing values were imputed via integration of peak areas.

Following recommendation guidelines for untargeted metabolomics^35^ metabolite features were retained when peaks were present in at least 70% of QC samples, relative standard deviation was less than 30%, and the extraction blank to mean QC peak area was less than 50%. Sample drift was corrected for using probabilistic quotient normalisation, then missing values were imputed using a Random Forest algorithm. Data were log transformed prior to analysis.

Raw spectral data files have been deposited to the MetaboLights^36^ repository with the study identifier MTBLS13077.

#### Statistical analysis

Unless otherwise indicated, data is presented as mean ± SD. Statistical analyses for respirometry, immunofluorescence, contractile function and Western blotting were performed using GraphPad Prism. Statistical analysis for transmission electron microscopy was performed using STATA. Contractile function was analysed using a two-way ANOVA (group x time). Immunofluorescence, respirometry, microscopy and Western blot images were analysed using a one way ANOVA. Post-hoc analysis was performed using Tukey’s range test. To assess sex differences in immunofluorescence, 95% confidence intervals for mean differences at 26 months were calculated and a two way ANOVA (age x sex) was performed. Statistical significance was set at *P* < 0.05.

Analysis of metabolomics data was performed using R. Differential metabolite abundance was determined using a linear model with empirical Bayes moderated t-tests. False discovery rate was accounted for using the Benjamini-Hochberg adjustment method and statistical significance was set at FDR adjusted *P* value < 0.05. Over representation analysis of significant metabolites was performed using MetaboAnalyst 5.0^37^.

## Results

### Ageing leads to increased total body weight and a reduction in muscle mass

To establish the effect of age on body composition, we assessed total body and tissue specific weights at each timepoint (Figure 1). Total body weight increased with age in male and female sedentary mice (Figure 1B). In contrast, quadriceps and gastrocnemius muscles significantly decreased from 3 to 24 and 26 months in both sexes (Figure 1D, 1E). Likewise, total lean mass decreased significantly from 3 to 26 months in sedentary females (Figure 1C).

**Figure 1.**
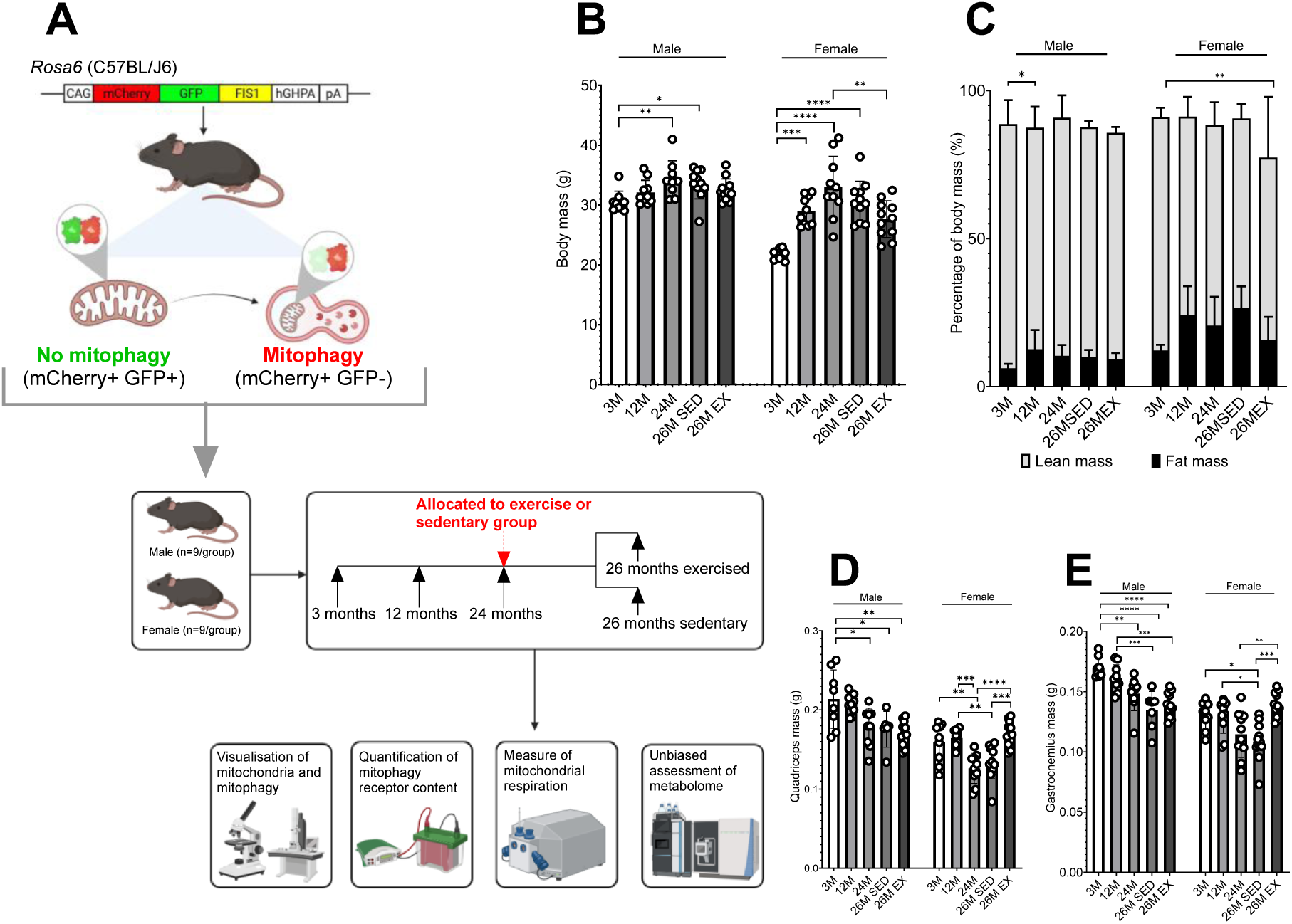
Age results in loss of muscle mass in male and female mice. (A) Experimental design studying mitophagy in young (3 months), middle-aged (12 months) and aged (24-26 months) mito-QC reporter mice. Created with BioRender.com. (B) Body mass (g) of male and female mito-QC reporter mice. (C) Proportion of fat and lean mass relative to total body mass (%) of male and female mito-QC reporter mice. (D) Mass (g) of quadriceps muscle in male and female mito-QC reporter mice. (E) Mass (g) of gastrocnemius muscle in male and female mito-QC reporter mice. Data are represented as mean ± SD (n=7-12). One-way ANOVA with multiple comparisons, *p<0.05, **p<0.01, ***p<0.001, ****p<0.0001.

We next examined the effect of exercise training and found significant increases from 24 to 26 months in quadriceps and gastrocnemius muscle mass in trained female mice only (Figure 1D, 1E). Trained female mice also had higher muscle mass at 26 months than their sedentary counterparts. Notably, male mice did not exhibit increased muscle mass following exercise training, indicating sex specific differences in the exercise response. As there were no significant differences between 24 and 26 months of age in sedentary mice, 12- and 24-month groups were excluded from further analysis.

### Skeletal muscle mitophagy increases with age in sedentary male mice

Immunofluorescence was used to visualise mitophagy in longitudinal sections of gastrocnemius muscle (Figure 2A, 2B, Extended Data Figure 2). Expression of GFP, used as a surrogate marker for mitochondrial content, was not affected by age in male sedentary mice (Figure 2C), however there was increased expression of mCherry from 3 months (p<0.001, Figure 2D). In contrast, female exhibited a concurrent increase in expression of GFP and mCherry (p<0.05 and p<0.01, respectively).

**Figure 2.**
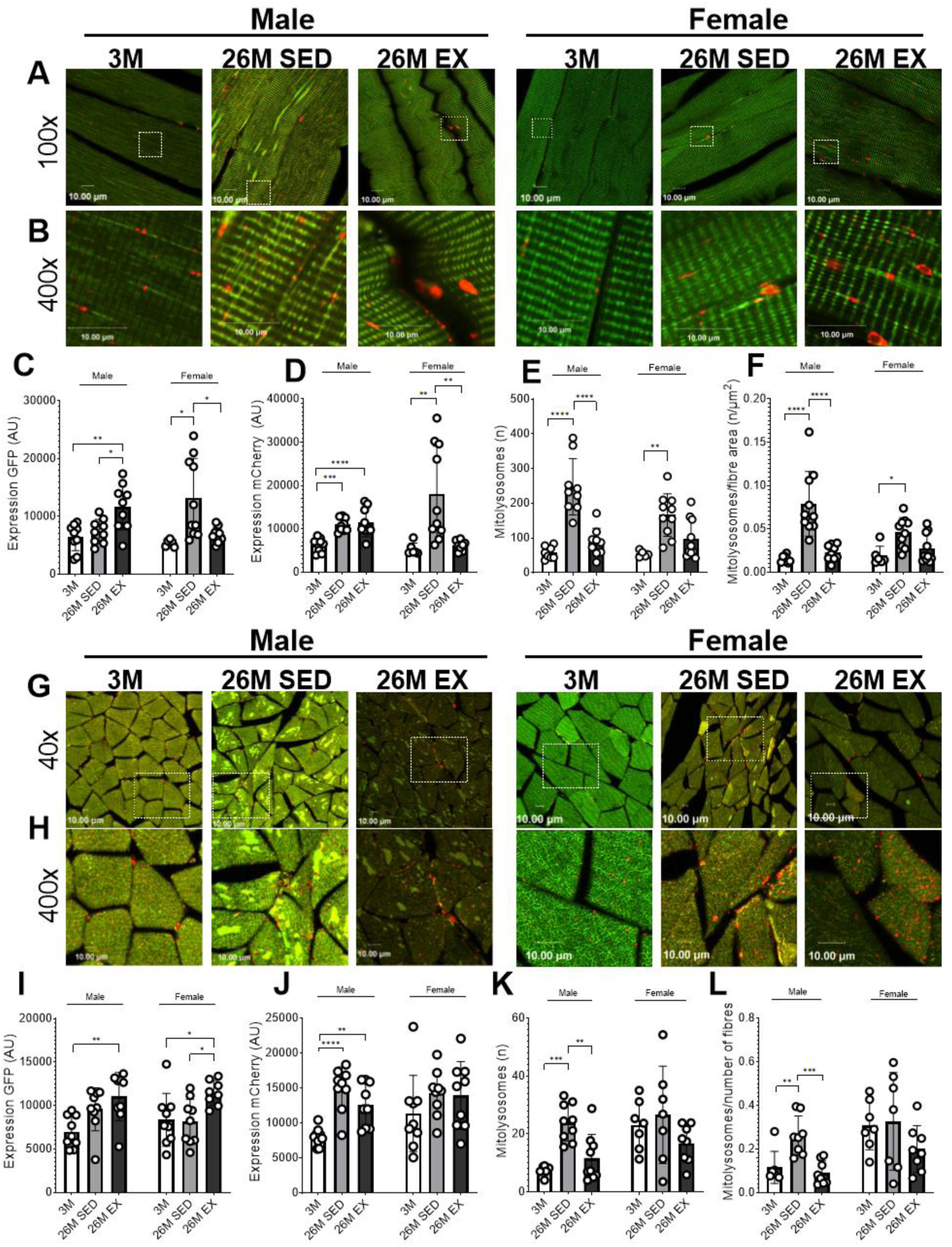
Exercise counteracts age related increases in mitophagy. (A, B) Representative confocal images of longitudinal gastrocnemius skeletal muscle fibres in male and female mito-QC reporter mice at 100x-(A) and 400x-(B) magnification. (C) Measure of GFP fluorescent intensity in fibres. (D) Measure of mCherry fluorescent intensity in fibres. (E) Number of mitolysosomes detected per image. (F) Number of mitolysosomes per fibre area (µm^2^) of image. (G, H) Representative confocal images of cross-sectional gastrocnemius skeletal muscle fibres in male and female mito-QC reporter mice at 40x-(G) and 400x (H) magnification. (I) Measure of GFP fluorescent intensity in fibres. (J) Measure of mCherry fluorescent intensity in fibres. (K) Number of mitolysosomes detected per image. (L) Number of mitolysosomes per number of fibres per image. Data represented as mean ± SD (n=6-10). One-way ANOVA with multiple comparisons, *p<0.05, **p<0.01, ***p<0.001, ****p<0.0001.

Given the magnitude of increase appeared greater in males than females, we examined whether there was a sex-specific effect on mitophagy (Extended Data Figure 3). Accordingly, number of mitolysosomes at 26 months was greater in sedentary males than females (p<0.05, 95% CI [12.25, 150.5]), which remained after normalising to fibre area (p<0.05, 95% CI [0.005, 0.06]). Additionally, significant age by sex interactions were observed when assessing number of mitolysosomes normalised to fibre area (p<0.05).

Composite immunofluorescent images of cross-sectional areas of gastrocnemius muscle were also acquired (Figure 2G, 2H, Extended Data Figure 4), which showed aged sedentary males had higher expression of mCherry (Figure 2J, p<0.0001) and more mitolysosomes (Figure 2K, p<0.001) than at 3 months. When mitolysosomes were normalised to number of fibres, the increase remained significant (Figure 2L, p<0.001). There was no change in mitolysosome number with age in female mice.

### Exercise training rejuvenates skeletal muscle mitophagy

We next examined the effect of exercise training on mitochondrial content. In males, there was a significant increase in expression of GFP in longitudinal (p<0.01, Figure 2C) but not cross-sectional fibres (Figure 2I). However, in both cases there was a significant increase in expression of mCherry (Figure 2D, 2J, p<0.0001 and p<0.01 for longitudinal and cross-sectional images, respectively). In males who underwent exercise training, mitolysosome number was not different from 3 months but was significantly reduced lower than their sedentary counterparts in longitudinal (Figure 2E, p<0.0001) and cross-sectional (Figure 2K, p<0.01) fibres, even when normalised to fibre area (Figure 2F, p<0.0001) or number of fibres (Figure 2L, p<0.001). Thus, eight weeks of exercise training prevented the age-related increase in mitophagy observed in male mice.

Female mice who underwent exercise training did not show any differences in expression of GFP, expression of mCherry or mitolysosome number from 3 months in longitudinal fibres. Expression of GFP increased from 3 months in cross-sectional fibres (Figure 2I, p<0.05), suggesting exercise training increased mitochondrial content in females.

### Accumulation of skeletal muscle mitochondrial aggregates occurs with age and are modulated by exercise

During acquisition of immunofluorescent images, we noted abnormal structures within muscle fibres which emitted GFP and mCherry fluorescence and were therefore deemed mitochondrial (Figure 3A). These structures, which we have termed aggregates, were observed in all groups except 3 month female mice. Aggregate size and number per fibre were recorded (Figure 3B-E). Although mean number of aggregates increased with age in sedentary males and females, the difference was not significant for either sex (p=0.09 and p=0.2, respectively). However, mean aggregate size did increase significantly from 3 to 26 months in sedentary mice (p<0.0001 and p<0.05 for males and females, respectively).

**Figure 3.**
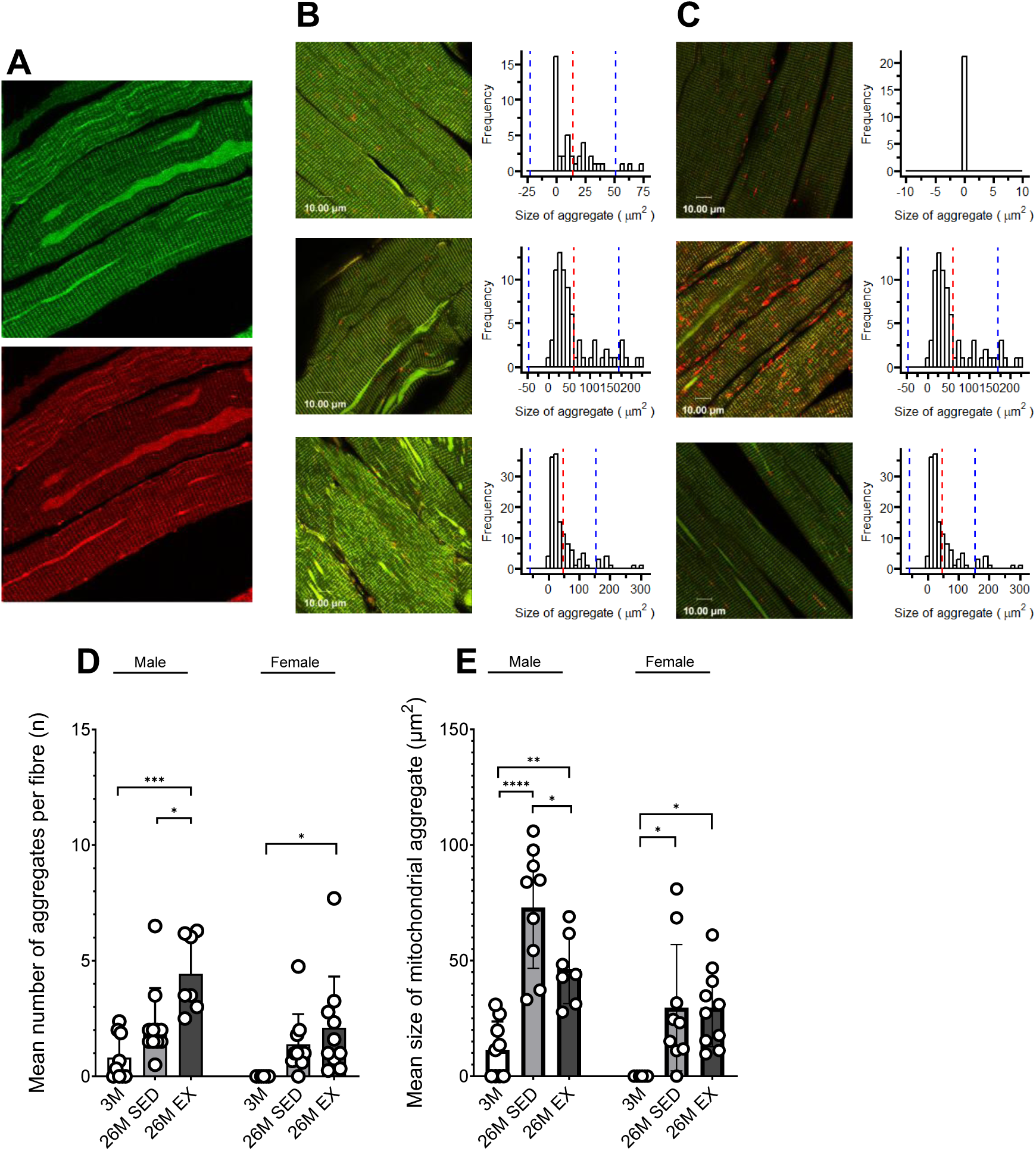
Ageing leads to accumulation of mitochondrial aggregates in muscle fibres. (A) Split channel fluorescence of representative images of aggregate features in a 26 month sedentary male mouse demonstrating the aggregates are mitochondrial structures. (B) Representative composite images of 3 month, 26 month sedentary and 26 month exercised male mice and distribution of aggregate size across ages. Light green areas were determined to be aggregates. (C) Representative composite images of 3 month, 26 month sedentary and 26 month exercised female mice and distribution of aggregate size across ages. (D) Mean number of aggregates per fibre in male and female mice. (E) Mean size of mitochondrial aggregates in male and female mice. Data represented as mean ± SD (n=6-9). One-way ANOVA with multiple comparisons, *p<0.05, **p<0.01, ***p<0.001, ****p<0.0001.

We found exercise training increased the mean number of aggregates from 3 months in males (p<0.001) and females (p<0.05), which suggested exercise does not prevent formation of aggregate structures. Aggregate size was unaffected by exercise in female mice, but old male mice who underwent exercise training had smaller aggregates than their sedentary counterparts (p<0.05).

### Skeletal muscle mitochondrial volume is increased by exercise training

Transmission electron microscopy was used to visualise whole mitochondria at 3 and 26 months (Figure 4A). Although total mitochondrial volume density was not significantly reduced with ageing in either sex (Figure 4G), we examined whether localisation of mitochondria influenced total density. While there was a more pronounced reduction in the SS region than the IMF region, reduction in SS mitochondrial content was not significant (p=0.07). Analysis of individual numerical density found no change in IMF (Figure 4D) or SS mitochondria (Figure 4I) in either sex, while analysis of individual profile size revealed that ageing led to a reduction in SS profile size in males only (p<0.001, Figure 4J). Mitochondrial cristae density was not affected by ageing (Figure 4F). A composite measure of mitochondrial volume density and cristae density, providing an estimate of cristae surface area per muscle volume, appeared to be reduced in ageing in males (Figure 4K).

**Figure 4.**
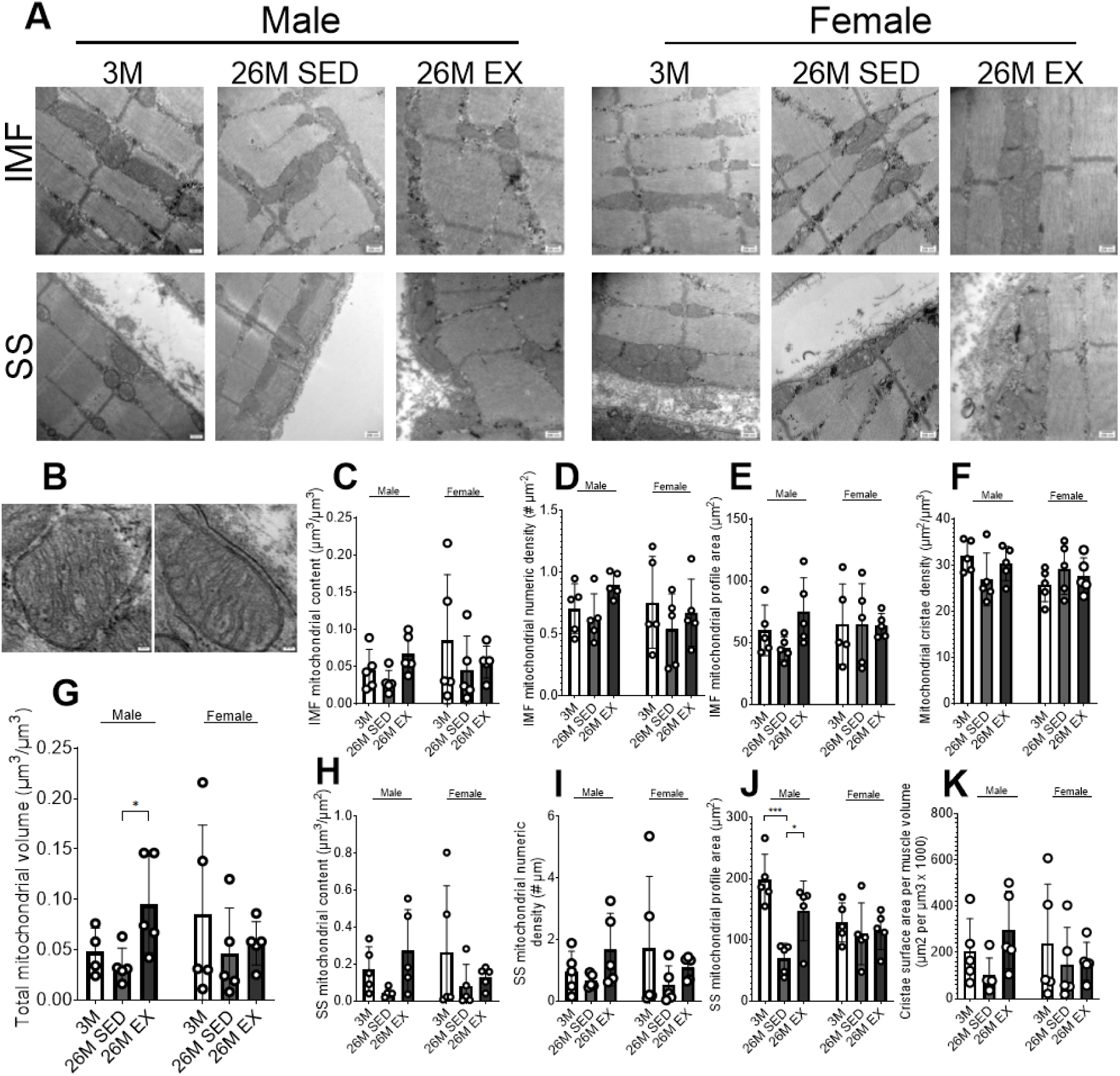
Mitochondrial volume decreases with age are prevented by exercise. (A) Representative transmission electron microscopy images of quadriceps muscle in male and female mice. Images were taken of intermyofibrillar (IMF) and subsarcolemmal (SS) mitochondria. (B) Representative image of IMF and SS mitochondria. (C) IMF mitochondrial content per muscle fibre (µm^3^/µm^3^). (D) Numeric density per image of IMF mitochondria (number mitochondria/µm^2^). (E) IMF mitochondrial profile area (µm^2^). (F) Mitochondrial cristae density. (G) Total mitochondrial volume (µm^3^/µm^3^) in skeletal muscle fibres. (H) SS mitochondrial content per fibre surface area (µm^3^/µm^2^). (I) Numeric density per image of SS mitochondria (number mitochondria/length of fibre (µm)). (J) SS mitochondrial profile area (µm^2^). (K) Composite measure of mitochondrial volume density and cristae density providing an estimate of cristae surface area per muscle volume (µm^2^/µm^3^ x 1000). Data represented as mean ± SD (n=5). One-way ANOVA with multiple comparisons, *p<0.05, ***p<0.001

Following exercise training, total mitochondrial volume density increased in males but not females (Figure 4G). This increase was ascribed to increased SS mitochondrial profile size, thus exercise training reversed the age-related reduction in mitochondrial profile size. Cristae density was not affected by exercise training. Cristae surface area per muscle volume tended to increase in males (p=0.09) but there was no difference in females.

### Exercise training restores skeletal muscle ADP sensitivity in aged mice

Mitochondrial respiration was assessed by high resolution respirometry (Figure 5A-G). There was no difference in oxygen flux in response to increased concentration of ADP with age in either male (Figure 5A) or female (Figure 5B) mice. Apparent Km significantly decreased with age in sedentary male mice (p<0.05, Figure 5C) but there was no change in female mice. Sex differences were also seen when assessing mitochondrial complex activity. Leak respiration through complex I was unaffected by age in sedentary males but significantly increased from 3 to 26 months in sedentary females (p<0.01, Figure 5D). Similarly, ADP stimulated respiration through complex I was significantly increased with age (p<0.05) in sedentary female mice while there was no difference in males (Figure 5E). ADP stimulated respiration through complexes I and II significantly decreased with age in sedentary males (p<0.01) but significantly increased in females (p<0.05, Figure 5F). Finally, maximal uncoupled respiration decreased significantly with age in sedentary males (p<0.05) but was unchanged in females (Figure 5G).

**Figure 5.**
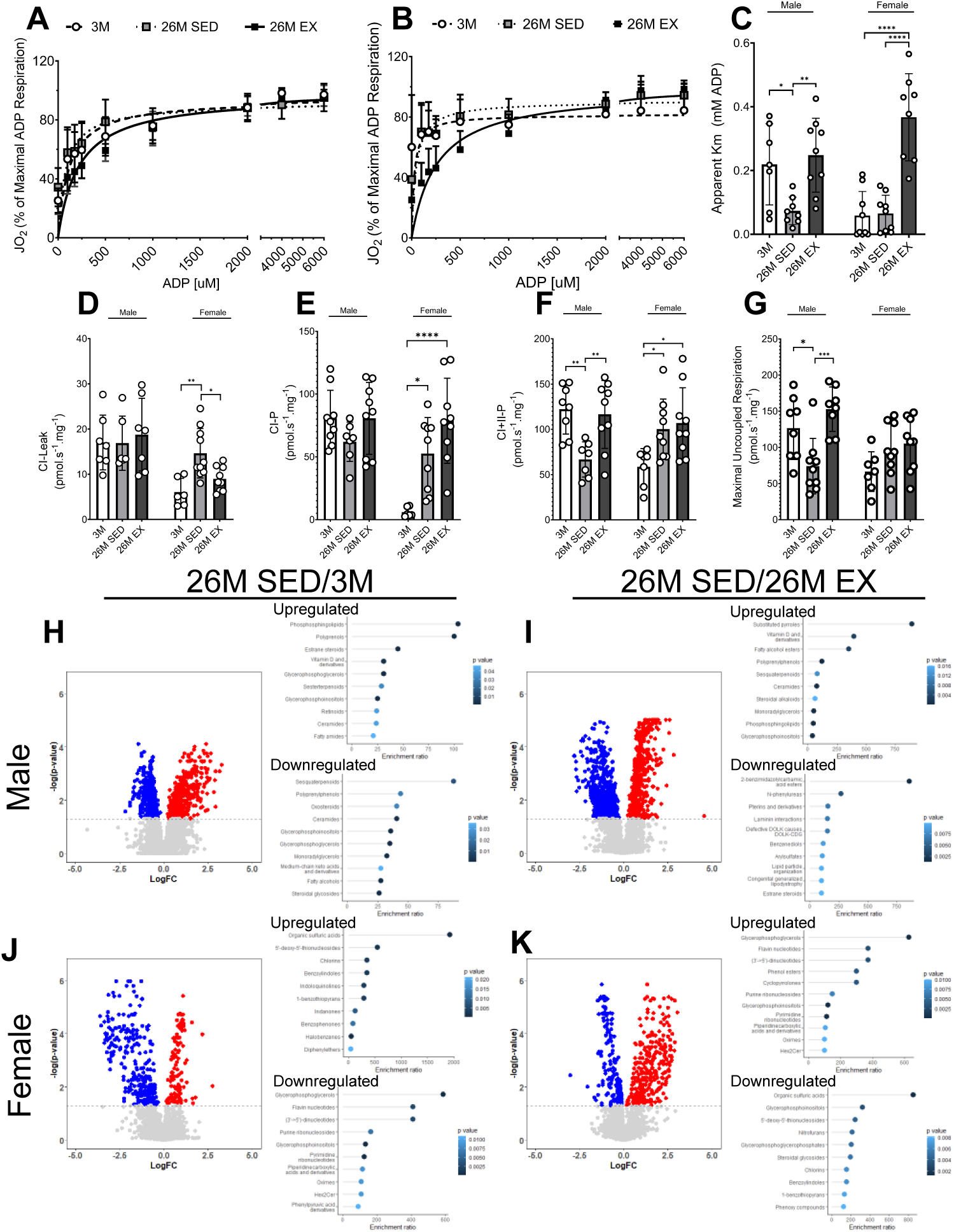
Exercise restores age-related loss of ADP sensitivity. (A, B) Oxygen flux in response to increasing concentration of ADP in 3 month, 26 month sedentary and 26 month exercised male (A) and female (B) mice. (C) Apparent Km for male and female mice. (D) Leak respiration through protein complex (C) I of the electron transfer system. (E) ADP stimulated respiration through CI of the electron transfer system. (F) ADP stimulation respiration through CI aznd CII of the electron transfer system. (G) Maximum uncoupled respiration in male and female mice. (H-K) Volcano plots of fold change versus p-value illustrating significant decreased (blue) or increased (red) metabolites in male (H, I) and female (J, K) mice where each point represents one metabolite, with lollipop plots showing enrichment ratio and p-value of top 10 enriched sub-classes of up- and downregulated metabolites. Respiration data represented as mean ± SD. One-way ANOVA with multiple comparisons (n=6-9), *p<0.05, **p<0.01, ***p<0.001

Following exercise training, oxygen flux increased (Figure 5A, B). Exercise training reversed the loss of apparent Km in male mice (Figure 5C) such that it was significantly greater than in age-matched sedentary counterparts (p<0.01). Apparent Km was also significantly greater following exercise training than at 3 (p<0.0001) or 26 (p<0.0001) months in female mice. Training did not change complex I leak or ADP stimulated respiration in males but complex I leak was significantly lower at 26 months in trained mice relative to sedentary counterparts (p<0.05, Figure 5D) and ADP stimulated respiration was significantly increased from 3 months in 26 month trained female mice (p<0.0001, Figure 5E). In males, ADP stimulated respiration through complexes I and II was significantly greater in trained mice than sedentary mice at 26 months (p<0.01) but there was no difference from 3 months (Figure 5F). Conversely, in females, respiration was significantly greater than 3 months following training (p<0.05) but was not different between sedentary and trained mice. Finally, maximal uncoupled respiration was significantly greater at 26 months in trained than sedentary male mice (p<0.001) but there had no effect in female mice (Figure 5G).

Having established that the effect of age on mitochondrial respiration is sex dependent and can be regulated by exercise, we next examined whether there were differential downstream metabolic effects. We examined quadriceps tissue using untargeted metabolomics and found large differences in the muscle metabolome between 3 and 26 months in both males and females (Figure 5H, J). In males, the most enriched subclasses in upregulated metabolites were phosphosphingolipids and polyprenols. The most highly enriched subclass of downregulated metabolites was sesquaterpenoids, followed by polyprenylphenols, and oxosteroids. Together, these datasets indicate large changes in lipid metabolism. In females, sulphuric acids were most highly enriched in upregulated metabolites while in downregulated metabolites glycerophosphoglycerols, flavin nucleotides, and dinucleotides were enriched, indicating changes to redox processes and DNA repair.

Exercise training also had potent effects on the muscle metabolome with male mice showing upregulation of vitamin D metabolites and fatty alcohol esters and downregulation of steroids and metabolites of benzenediols (Figure 5I). Female mice showed a different metabolic response, with the metabolites which were downregulated in sedentary mice being upregulated following exercise training (Figure 5K). Classes of upregulated metabolites included glycerophosphoinositols and glycerophosphates, suggesting exercise training had induced a change in membrane structure and cellular regulation.

### Exercise training improves skeletal muscle functional capacity

EDL and SOL muscles were tested by excitation contraction coupling to assess whether increased mitophagy was associated with reduced muscle function in older mice. In sedentary males, specific force output was significantly lower in 26 month mice compared to 3 month mice in both EDL (p=0.0281) and SOL (p=0.0403) muscles (Figure 6A, E). Sedentary old male mice also showed a greater rate of fatigue (p<0.0001) in EDL muscle relative to 3 month mice (Figure 6B), although there was no difference in fatiguability in SOL muscle (Figure 6C) and there was no difference in recovery between age groups. Sedentary older female mice showed no difference from 3 months in specific force (Figure 6C) or fatigue (Figure 6D) in EDL, but in the SOL specific force was significantly reduced (p<0.0001, Figure 6G) and older mice showed greater fatigue (p<0.0001) with poorer recovery (p<0.0001, Figure 6H).

**Figure 6.**
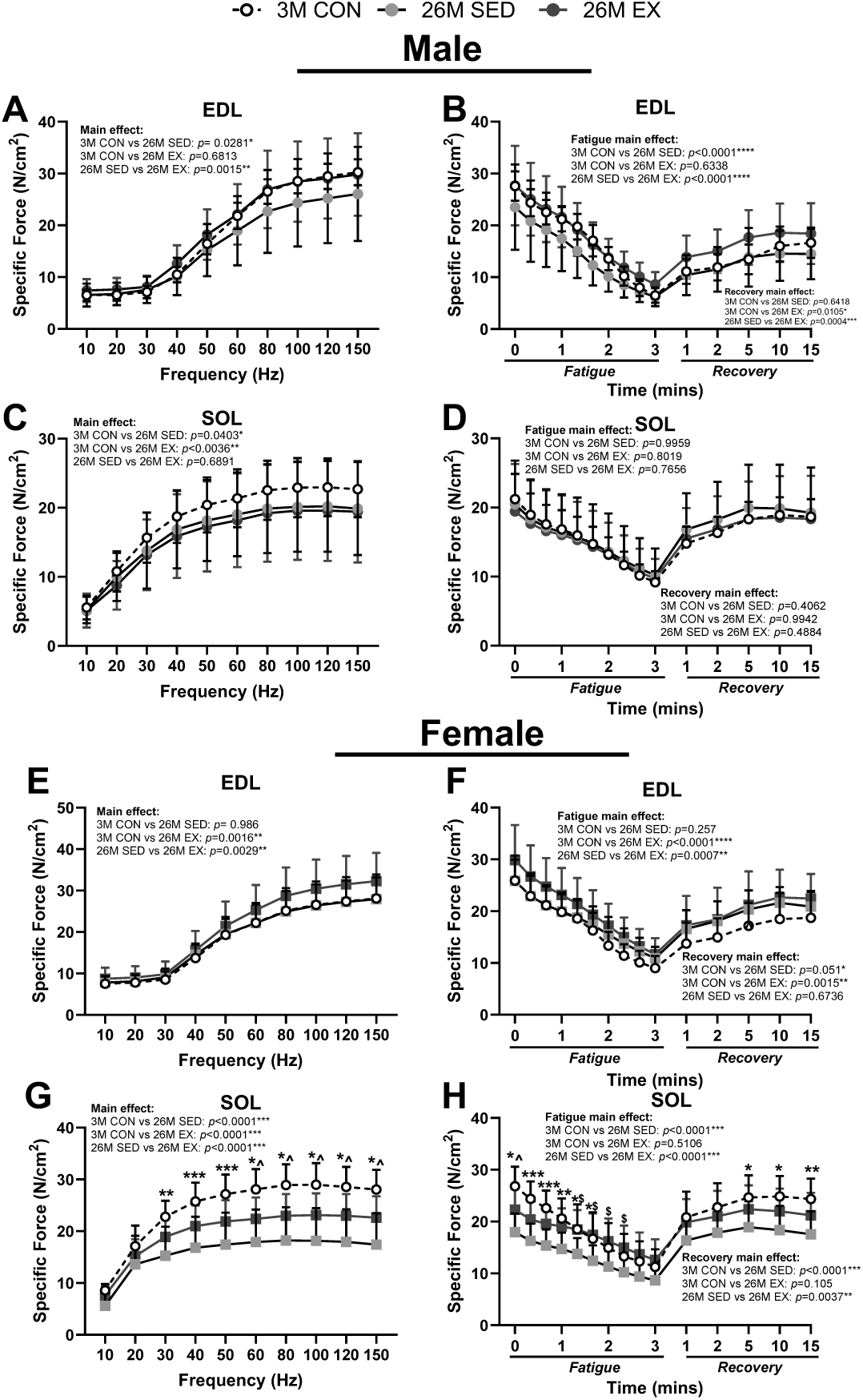
Exercise training improves muscle strength and fatigue resistance. (A) Force output normalised to muscle cross sectional area in male EDL muscle. (B) Force output normalised to muscle cross sectional area over a 3 minute fatiguing period and 15 minute recovery period in male EDL muscle. (C) Force output normalised to muscle cross sectional area in male SOL muscle. (D) Force output normalised to muscle cross sectional area over a 3 minute fatiguing period and 15 minute recovery period in male SOL muscle. (E) Force output normalised to muscle cross sectional area in female EDL muscle. (F) Force output normalised to muscle cross sectional area over a 3 minute fatiguing period and 15 minute recovery period in female EDL muscle. (G) Force output normalised to muscle cross sectional area in female SOL muscle. (H) Force output normalised to muscle cross sectional area over a 3 minute fatiguing period and 15 minute recovery period in female SOL muscle. Data represented as mean ± SD (n=8-10). Two-way ANOVA, *p<0.05, **p<0.01, ***p<0.001, ****p<0.0001; ^p<0.05 (3M CON vs 26M EX); $p<0.05 (26M SED vs 26M EX)

Following exercise training, 26 month male and female mice showed significantly greater specific force output than their sedentary counterparts in EDL muscle (male: p=0.0015, female: p=0.0029, Figure 6A, C) and had lower fatigue (male p<0.0001, female: p=0.0007, Figure 6B, D). Recovery was also significantly greater in exercise trained males (p=0.0004) but was not different from sedentary in females (Figure 5B, D). In SOL muscle, there was no difference in specific force output (Figure 6E) or fatigue and recovery (Figure 6F) between sedentary and exercise trained male mice at 26 months, but female mice had a significantly greater force output (p<0.0001, Figure 6G), lower fatigue (p<0.0001) and greater recovery (p=0.0037) following exercise training (Figure 6H).

### Basal mitophagy receptor content is increased by exercise training in male and female skeletal muscle

Given the alterations in basal mitophagy observed with age and following exercise training, we measured mitophagy receptor content in skeletal muscle. In sedentary male mice, age did not change the content of any receptor, and in sedentary females only BNIP3 content was significantly different from 3 to 26 months (p<0.001, Figure 7B). However, several changes were seen following exercise training. BNIP3 content increased significantly from 3 months (p<0.01) and between age matched male sedentary and trained mice (p<0.001) and was significantly greater from 3 months after exercise training (p<0.01) in females (Figure 7B). Male mice also exhibited significant increases between sedentary, and exercise trained mice in FUNDC1 (p<0.05, Figure 7C) and BCL2L13 (p<0.01, Figure 7D) which were not observed in females. FUNDC1 and BCL2L13 protein content was also significantly greater than at 3 months following exercise training (p<0.05 and p<0.01, respectively). In females, optineurin content was significantly increased (p<0.05) in trained mice relative to sedentary mice at 26 months (Figure 7E) while NIX content was significantly increased from 3 months following training (p<0.01, Figure 7F).

**Figure 7.**
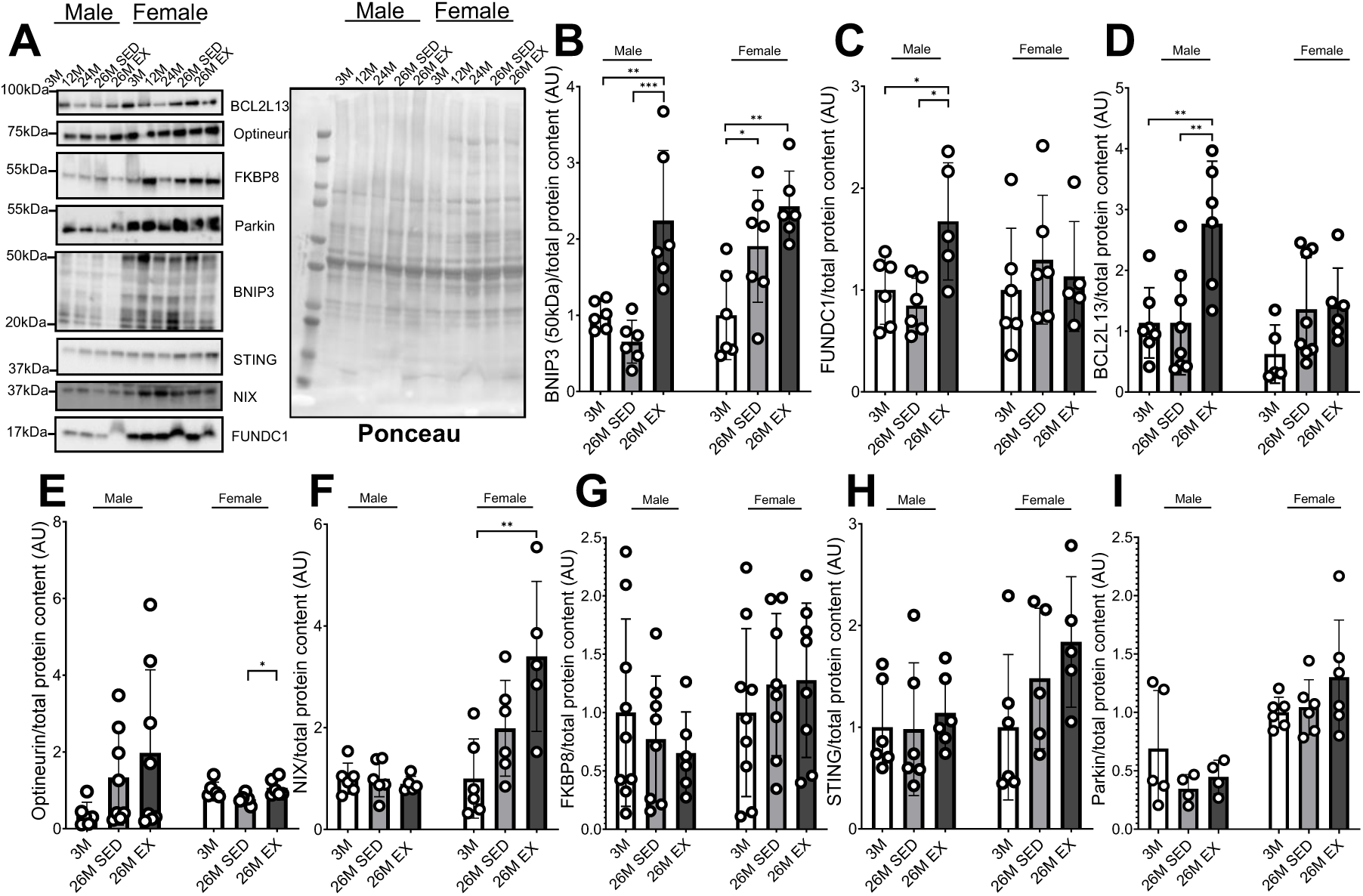
Exercise training increases basal expression of BNIP3, FUNDC1 and BLC2L13. (A) Representative Western blot showing total protein content visualised using Ponceau staining and representative bands for each receptor quantified. (B-I) Quantification (in arbitrary units) of fold change from 3 months for (B) BNIP3, (C) FUNDC1, (D) BCL2L13, (E) Optineurin, (F) NIX), (G) FKBP8, (H) STING, (I) Parkin. Band intensities were normalised to total protein content. Data represented as mean ± SD. One-way ANOVA with multiple comparisons (n=6-8), *p<0.05, **p<0.01, ***p<0.001

## Discussion

Our results clearly show that the development of sarcopenia occurs in parallel to a large increase in basal mitophagy in skeletal muscle of old, sedentary mice. The hyper-mitophagy response observed with age was completely reversed by 8 weeks of exercise training, providing a novel mechanism by which endurance exercise training improves skeletal muscle mitochondrial homeostasis. Our analysis also identified clear sex differences throughout the study which indicate mitophagy is increased to a greater extent in males and suggest a protective mechanism against loss of mitochondrial and muscle function exists in female mice at 26 months.

Sexual dimorphism related to mitochondria has been reported in previous studies. Mitochondrial content has been shown to be reduced with age in male but not female cohorts^38^ and PINK1, a kinase involved in stress-induced mitophagy, had a 75% higher expression in old male than female rats^39^. This study provides further evidence showing sex differences exist in mitochondrial quality control mechanisms and may underlie differential adaptations in respiratory capacity muscle function and sarcopenia progression. It is possible that the onset of mitochondrial dysfunction would occur at an older age in female mice, as sarcopenia progression typically occurs later in female mice. However we instituted a cutoff of 26 months to prevent the occurrence of comorbidities which are common in mice over the age of 2 years^40,41^. In doing so, we can attribute age as the sole driver behind the observed changes in mitophagy.

Utilising mito-QC mice, we revealed an accumulation of mitochondrial aggregate structures within skeletal muscle fibres that occurred with age in both sexes. Similar structures have been noted in previous work^19^, although they have not yet been studied in detail. We defined a set of criteria to appropriately quantify the structures and found aggregate size was linked to both age and activity level in male mice, aligning with the observed hyper activation of mitophagy. Further, as exercise restored levels of mitophagy comparable to 3 month old mice in skeletal muscle, the size of the mitochondrial aggregates was also reduced. Why mitochondrial aggregates are formed is currently unclear, however the appearance of giant mitochondrial structures with age has been observed before and linked to hyper fusion events, thought to be triggered in an attempt to rescue mitochondrial dysfunction^42^.

Whilst reduced mitochondrial content is common with increased age^1^, our data suggests that localisation also dictates this process. We observed reduced SS profile area with increased age suggesting that SS mitochondria were more susceptible to loss than IMF mitochondria in male mice. Subpopulations of mitochondria are known to play distinct roles in metabolism^43^ and have demonstrated independent adaptations to exercise training interventions^44^. Multiple studies have shown the protein content of IMF mitochondria is higher than that of SS mitochondria and IMF mitochondria have greater enzyme activity than SS mitochondria^45–48^. Specifically, IMF mitochondria express higher levels of proteins associated with oxidative phosphorylation and a higher respiratory chain complex^49^. IMF mitochondria, which are thought to be more important for energy production in contractile activity, may therefore be preferentially retained in periods of increased mitophagy. In contrast, exercise increased mitochondrial volume in male mice, in keeping with previous reports^50^, which we attribute to increased SS profile area. Given the clear reversal of the detrimental effect of age on SS profile area following training, we suggest that regulation of mitophagy through exercise training reverses the effect of age on mitochondrial availability by inducing proliferation of mitochondrial populations which were preferentially lost in sedentary mice.

In addition to analysing the ultrastructure of mitochondria we assessed functional capacity. It is well documented that age can blunt mitochondrial ADP sensitivity^51^ and that individuals who have lower physical activity report low K_m_ values, while highly active individuals report high K_m_ values^52^. It is also recognised that exercise is a potent regulator of mitochondrial ADP sensitivity^53^. In the present study, mitochondrial sensitivity was reduced with age in sedentary old male, but not female mice, aligning with the observed increase in mitophagy. Furthermore, exercise, which returned basal mitophagy at 26 months to levels comparable with young mice, was able to restore sensitivity to baseline in males. Thus, we suggest the hyper-activation of mitophagy is responsible for age-related losses of ADP sensitivity and that exercise regulates ADP sensitivity through its control of mitophagy.

To evaluate wider effects of age and increased mitophagy, we adopted an unbiased approach to identify differences in the metabolome between 3 month and 26-month sedentary mice. Distinct differences in up- and downregulated metabolites were observed between the sexes. Lipid metabolism was strongly affected by age in sedentary male mice, but not female mice. For instance, two classes of sphingolipids, ceramides and their downstream metabolites phosphosphingolipids, were upregulated with age. Sphingolipids have been repeatedly shown to accumulate with age^54,55^, which has been suggested as a driver of mitophagy^56^ due to their pro-inflammatory properties^57^. The observed upregulation may therefore be promoting an inflammatory environment that contributes to hyperactivation of mitophagy in aged mice. Other upregulated lipid classes included polyprenols, which are known to accumulate with age in other tissues including spleen^58^ and brain^59^ and have antioxidant properties. As oxidative stress is a known trigger of mitochondrial damage and given the loss of mitochondrial content and function in sedentary males, upregulation of polyprenols is likely a protective mechanism to prevent further mitochondrial dysfunction. Downregulated lipids were found to have roles in the electron transport chain^60^, which is likely linked to the reduction in ADP sensitivity previously observed. Beyond lipids, oxosteroids were found to be downregulated in sedentary males. Androgen deficiency in older males has previously been linked to decreased muscle mass^61,62^, aligning with measurements of total lean mass and mass of exercised skeletal muscle of mice in this study.

In contrast, metabolites upregulated in female mice were sulphuric acid metabolites, indicative of increased xenobiotic metabolism. Downregulated metabolites were found to be involved in energy metabolism, notably playing key roles in the transfer of electrons during oxidative phosphorylation^63^ and in the production of ATP^64^. This metabolomic profiling suggests that while mitochondrial sensitivity is not reduced in older females, there are still effects of ageing on energy metabolism, suggesting that mitochondrial dysfunction occurs at an older age in females. Interestingly, the potential negative effects on energy metabolism with age were reversed by exercise, as the same metabolite classes were upregulated in trained mice, further demonstrating the beneficial effects of exercise training on aged mice.

Reduced muscle strength is a common comorbidity of ageing and it has been repeatedly shown that low muscle strength in older adults is dependent on activity levels^65–68^. Our data support this, showing a reduced force output of EDL muscle from sedentary old male mice, but no difference in specific force output of EDL muscle between young and old exercise trained male mice. Sedentary old male mice also fatigued easier than both young and exercise trained old mice, and had a blunted recovery compared to exercise trained mice. In females, there was no difference in specific force output of EDL muscle between young and sedentary old mice. There was also no difference in fatigue between 3 month and 26 month sedentary groups, although interestingly, recovery was better in older mice than at 3 months. SOL muscle was weaker at 26 months than baseline regardless of activity in both sexes. Despite having a lower force output, female mice who underwent exercise training were more fatigue resistant and had better recovery than sedentary old mice, which was not observed in male mice. In addition to a greater loss of strength, we found loss of muscle mass was exacerbated in male mice when compared to female mice, showing a clear association between mass and function, symptomatic of sarcopenia^69^.

Finally, to understand the mechanistic basis of the hyper-mitophagy response observed, we quantified basal content of known mitophagy receptors. No change in protein content could be linked to hyper-mitophagy. However, BNIP3, FUNDC1 and BCL2L13 content increased after exercise training. BNIP3 and FUNDC1 both regulate mitophagy in response to hypoxia^70,71^. In addition, FUNDC1 is an essential regulator of mitochondrial fission and fusion through its interaction with LC3^72^. BCL2L13 induces mitophagy through the recruitment of the ULK1 complex^73^ and also regulates calcium signalling at mitochondria-endoplasmic reticulum contact sites^74^. Exercise is a potent stimulator of mitophagy receptor content, and BNIP3, FUNDC1 and BCL2L13 have all previously been reported to increase in skeletal muscle following chronic exercise training^75–77^. Thus, the reduction of mitophagy following exercise training in aged mice may have been driven by the upregulation of mitophagy receptors to increase basal mitophagy. Alternatively, exercise may have led to an increase in lysosomal biogenesis to facilitate greater mitochondrial clearance, given the established links between exercise and lysosomal biogenesis^78^.

In summary, our study is the first to report that sarcopenia is characterised by hyper-mitophagy in skeletal muscle, a process that is reversed by exercise training. High resolution microscopy demonstrated that localisation is an essential component in dictating loss of mitochondrial volume in sarcopenia. Additionally, blunted mitochondrial function in sarcopenic mice can be reversed by exercise training. Metabolomics indicated large changes to lipid metabolism are characteristic of sarcopenia, whereas exercise has a beneficial effect on membrane structure and cellular regeneration. Importantly, exercise is capable of reversing the negative effects of sarcopenia on energy metabolism. Finally, while sarcopenia had no effect on mitophagy receptor content, exercise training appears to increase basal content of FUNDC1, BNIP3 and BCL2L13. Collectively our data supports a role for mitophagy in the progression of sarcopenia and suggests that therapeutic targeting of mitophagy may be a novel approach to target sarcopenia via an improvement in mitochondrial quality control.

## Acknowledgements

IA and AP were supported by Wellcome Leap’s Dynamic Resilience Program (jointly funded by Temasek Trust). This research was facilitated by access to Sydney Mass Spectrometry, a core research facility at the University of Sydney. The authors acknowledge the technical and scientific assistance of Sydney Microscopy & Microanalysis, the University of Sydney node of Microscopy Australia.

## Supplementary methods

### Optimisation of Leica SP8 confocal microscope laser power and settings for mitolysosome detection

Laser power was adjusted until exposure reached submaximal saturation (i.e. when blue pixels began to appear within the tissue) for both GFP and mCherry autofluorescence. Overexposure was performed on three mice and mean optimal laser power was calculated for each protein. mCherry fluorescence was set to 552nm. GFP fluorescence was set to 488nm. Line averaging and pixel resolution settings were optimised on longitudinal samples collected from 3 month male mice (Supplementary Figure 2). Increasing pixel resolution did not significantly improve number of mitolysosomes detected but did increase time taken to capture images. As such, pixel resolution was set at 512×512. Photo bleaching occurred at line averages 16 and 32, therefore line averaging was set at 4.

### Optimisation of mito-QC Counter macro settings for detection of mitoQC skeletal muscle tissue

The *mito-QC Counter* macro was designed for use in mitoQC cell lines and required optimisation for skeletal muscle tissue. Number of mitolysosomes in representative images was counted by eye and recorded using the ImageJ multi-point clicker function to establish an accurate control. A second arbitrary control was established by using the ImageJ particle counter to detect number of mitolysosomes. The *mito-QC Counter* macro contains three primary settings that can be optimised by the user: the ratio threshold, which instructs the macro to exclude pixels below a certain ratio of mCherry to GFP; smoothing radius, the value that instructs the macro to distinguish high ratio pixels that are in close proximity to each other; and the standard deviations (SD) above red mean, which instructs the macro to only consider pixels above a determined threshold defined by each image’s mean mCherry intensity.

To optimise ratio threshold, the setting was incrementally increased from the default setting of 0.5 AU and number of mitolysosomes detected was compared to the number counted by eye. The most accurate quantification of mitophagy compared to by eye measurements was found to be 0.8 AU (Supplementary Figure 3). For all further analysis, ratio threshold was set at 0.8 AU. The same procedure was followed to optimise smoothing radius, which began at the default setting of 1. Increased smoothing radius did not improve accuracy of mitolysosome detection, therefore smoothing radius was set at 1 (Supplementary Figure 4). Laser power settings of the SP8 confocal microscope had previously been optimised for a maximal fluorescence intensity of 20,000 when imaging mitoQC tissues as both GFP and mCherry fluorescence intensities peaked at 10,000. SD above red mean was therefore normalised to a fluorescence intensity of 10,000 for all analysis.

---

**Supplementary Figure 1.**
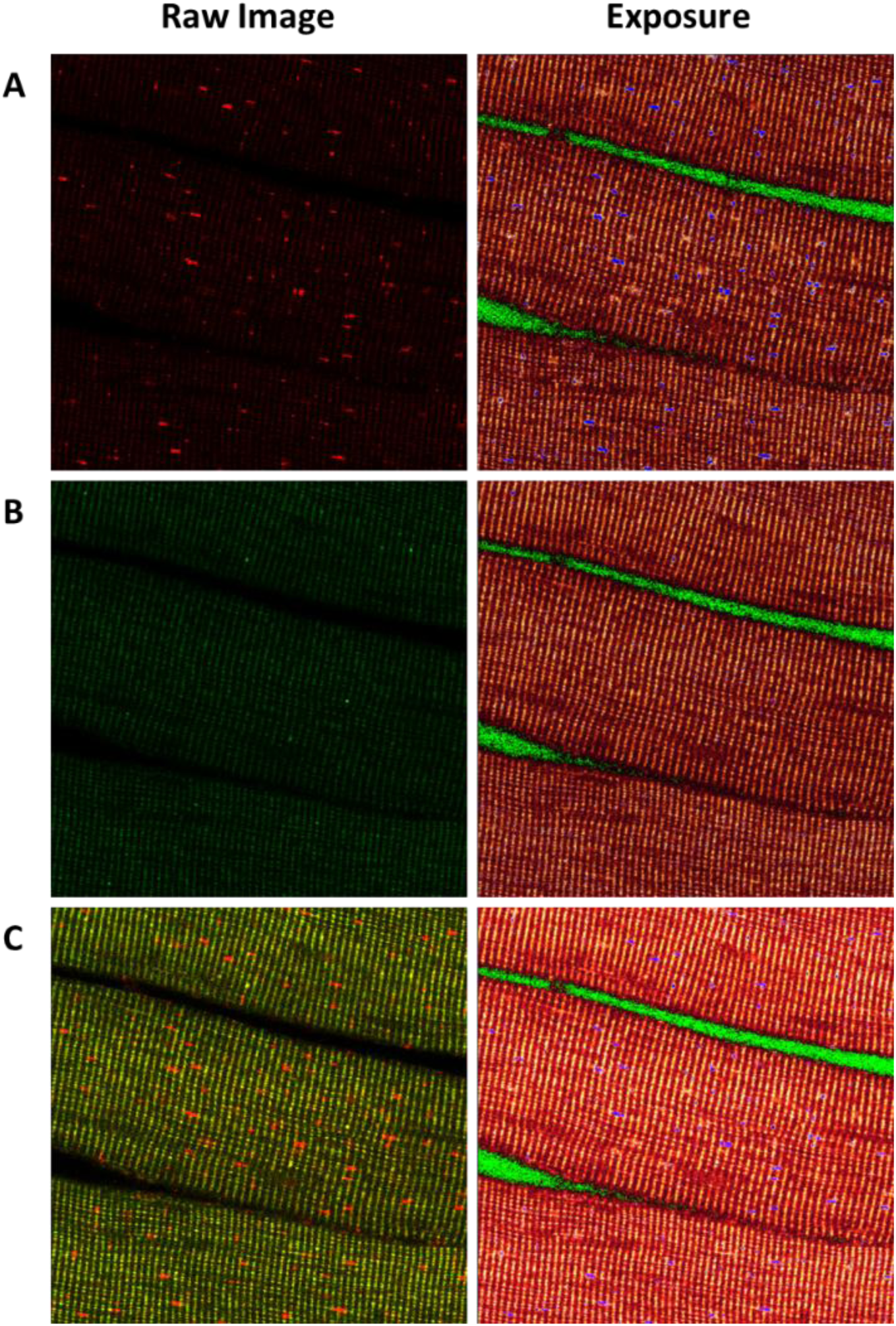
Optimisation of confocal microscope settings for skeletal muscle from mitoQC mice. (A) Optimised mCherry fluorescence of gastrocnemius tissue from 12-week mitoQC male mice and its corresponding exposure under confocal microscope laser at 552nm. (B) Optimised GFP fluorescence of gastrocnemius tissue from 12-week mitoQC male mice and its corresponding exposure under confocal microscope at 488nm. (C) Composite image of mCherry and GFP fluorescence and its corresponding exposure.

**Supplementary Figure 2.**
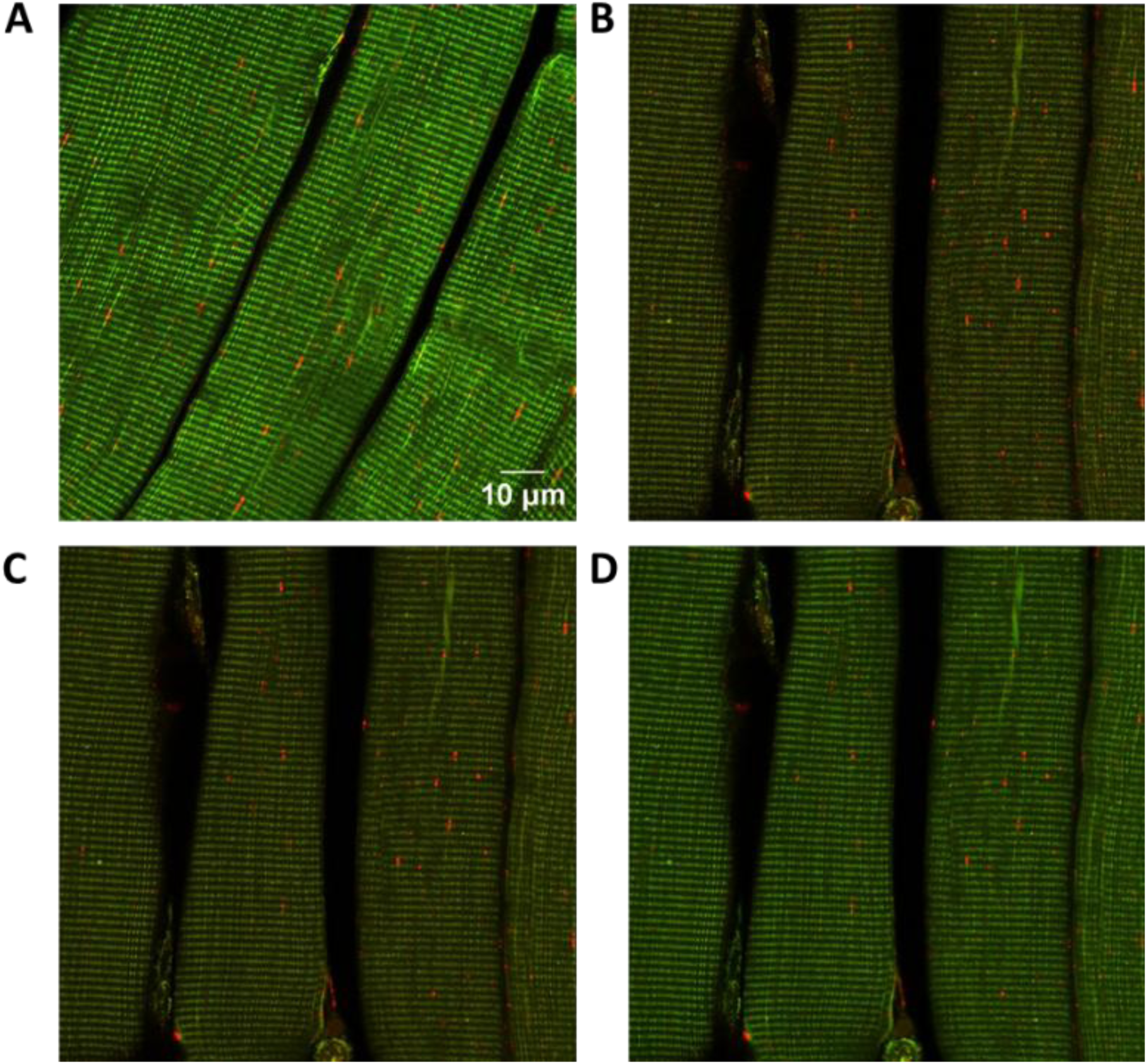
Optimisation of pixel and line average settings for skeletal muscle from mitoQC mice. (A) Representative image of longitudinal gastrocnemius muscle at 1024×1024 pixels with line average 64. (B) Representative image of longitudinal gastrocnemius muscle at 512×512 pixels with line average 4. (C) Representative image of longitudinal gastrocnemius muscle at 512×512 pixels with line average 16. (D) Representative image of longitudinal gastrocnemius muscle at 512×512 pixels with line average 32

**Supplementary Figure 3.**
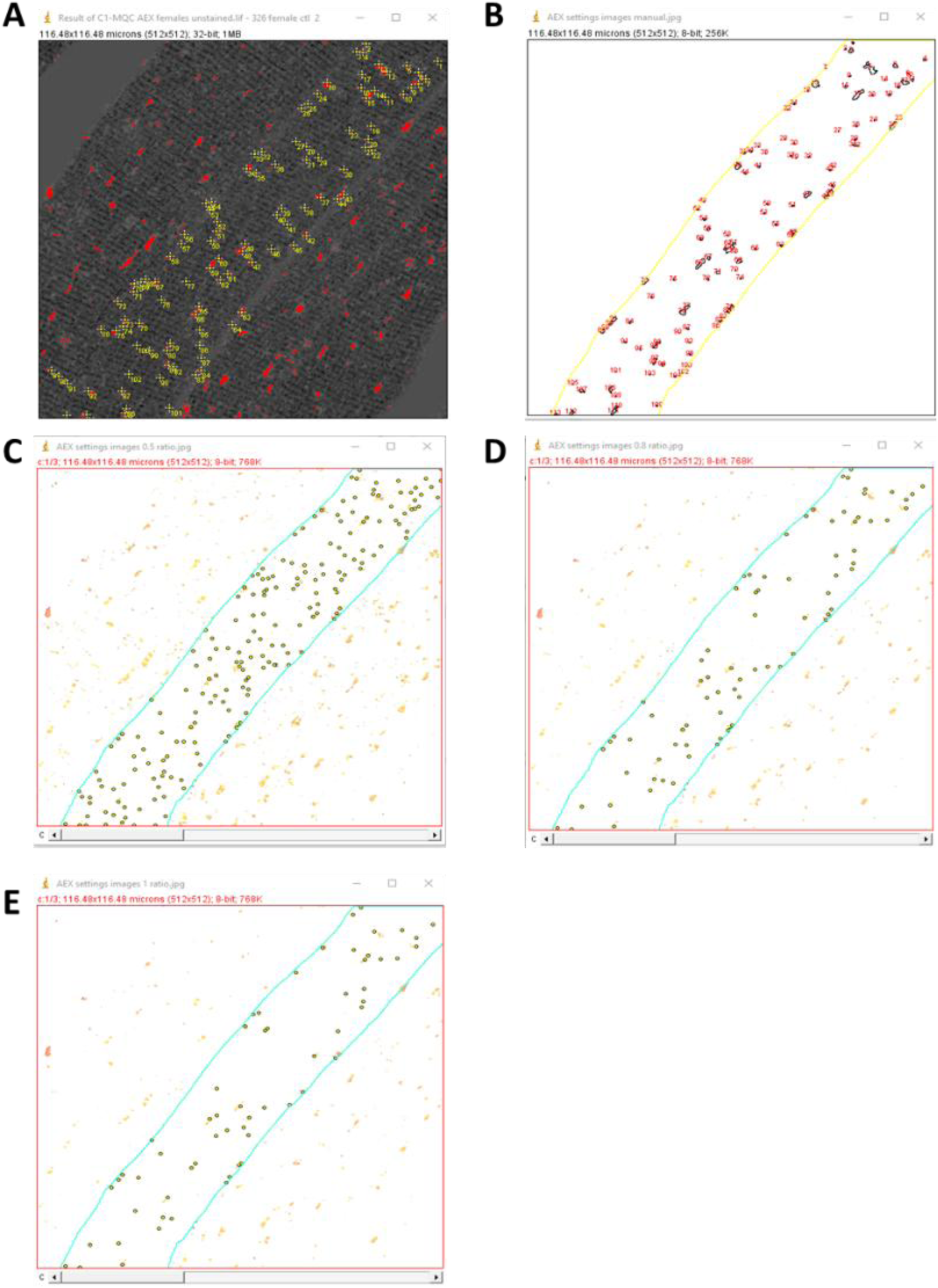
Optimisation of ratio threshold. (A) By eye count of mitolysosomes in gastrocnemius muscle from 12-week mitoQC female mice. (B) Skeletonised muscle fibre showing mitolysosomes detected by ImageJ particle counter plugin. (C-E) Skeletonised muscle fibre showing mitolysosomes detected by *mito-QC Counter* macro at ratio threshold (C) 0.5, (D) 0.8, (E) 1.

**Supplementary Figure 4.**
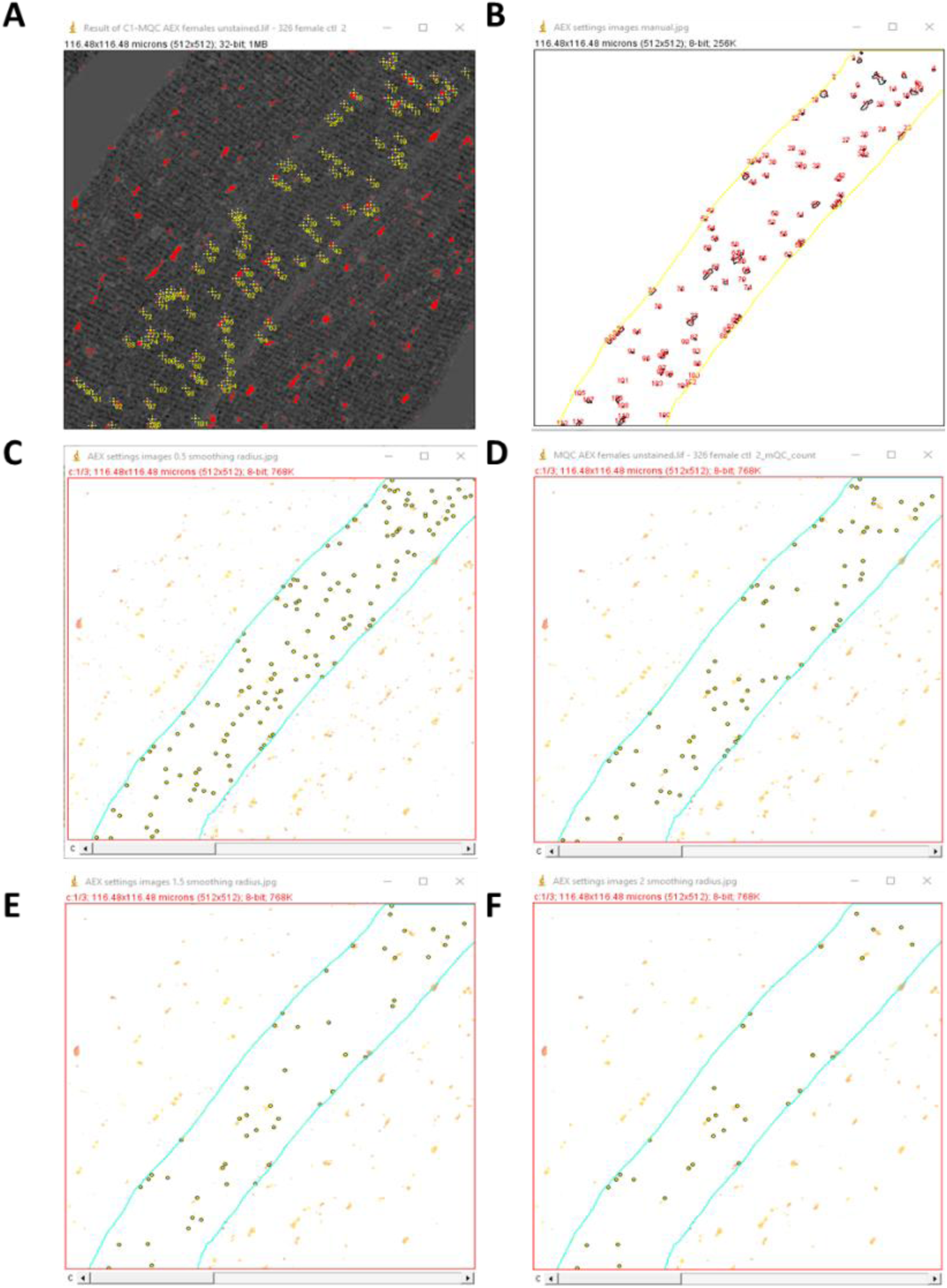
Optimisation of smoothing radius. (A) By eye count of mitolysosomes in gastrocnemius muscle from 12-week mitoQC female mice. (B) Skeletonised muscle fibre showing mitolysosomes detected by ImageJ particle counter plugin. (C-F) Skeletonised muscle fibre showing mitolysosomes detected by *mito-QC Counter* macro at smoothing radius (C) 0.5, (D) 1, (E) 1.5, (F) 2.

**Extended data figure 1.**
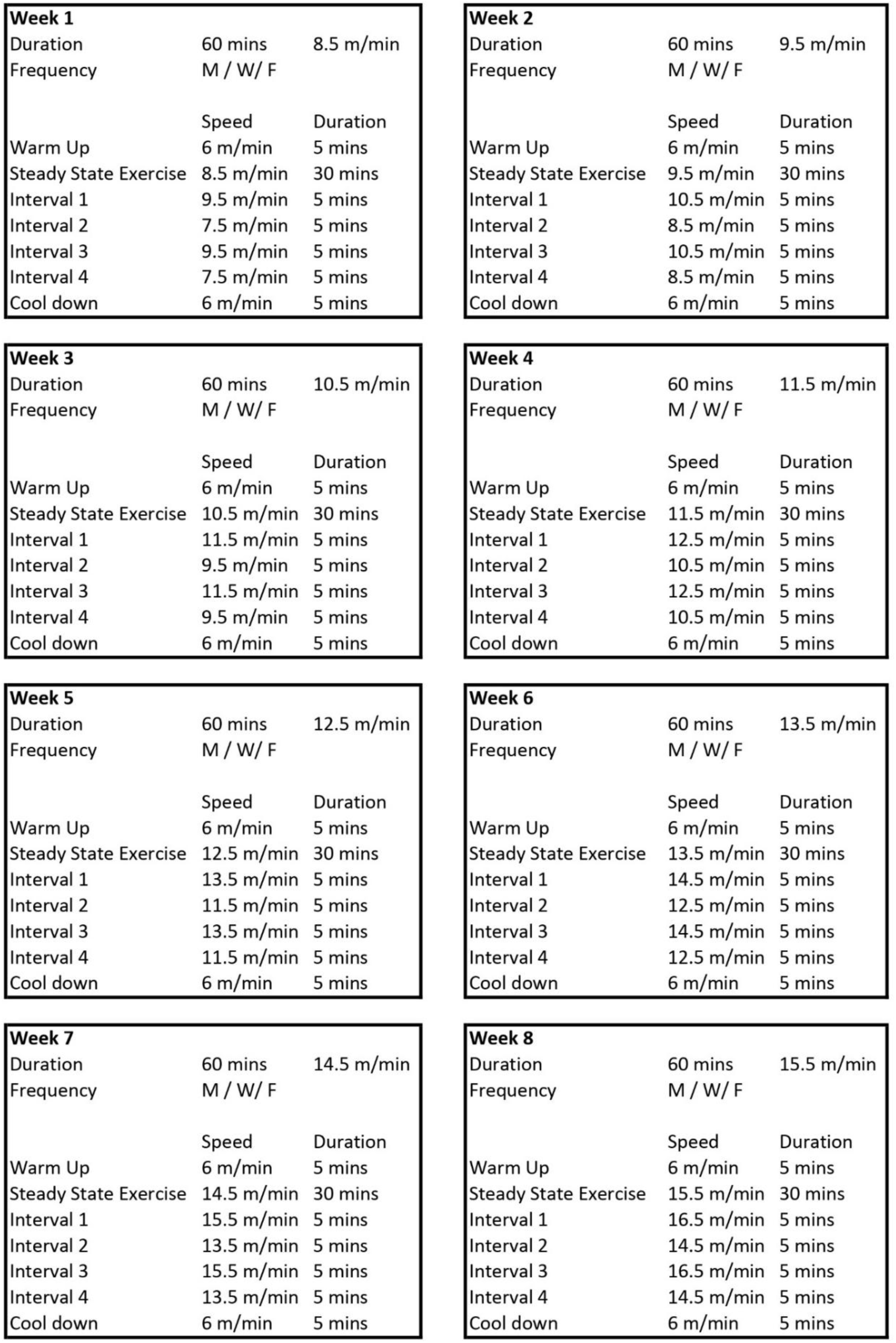
**8 week progressive endurance training protocol for male and female mice at 24 months**

**Extended data figure 2.**
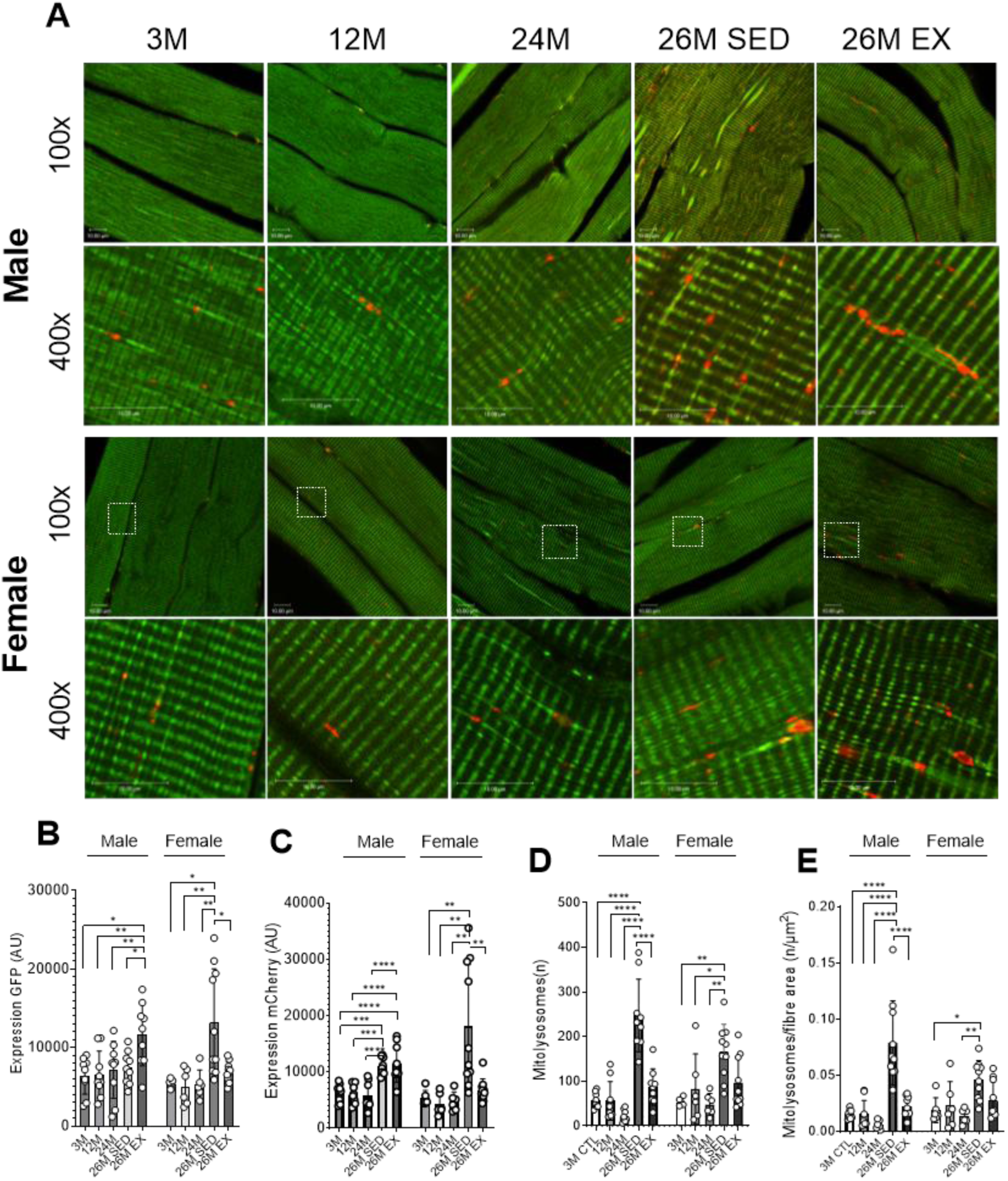
Immunofluorescent imaging of longitudinal sections of gastrocnemius muscle from 3 to 26 months. (A) Representative confocal images of longitudinal skeletal muscle fibres in male and female mice at 100x- and 400x- magnification. (B) Measure of GFP fluorescent intensity in fibres. (C) Measure of mCherry fluorescent intensity in fibres. (D) Number of mitolysosomes detected per image. (E) Number of mitolysosomes per fibre area (µm^2^) of image. Data represented as mean ± SD (n=6-10). One-way ANOVA with multiple comparisons, *p<0.05, **p<0.01, ***p<0.001, ****p<0.0001

**Extended data figure 3.**
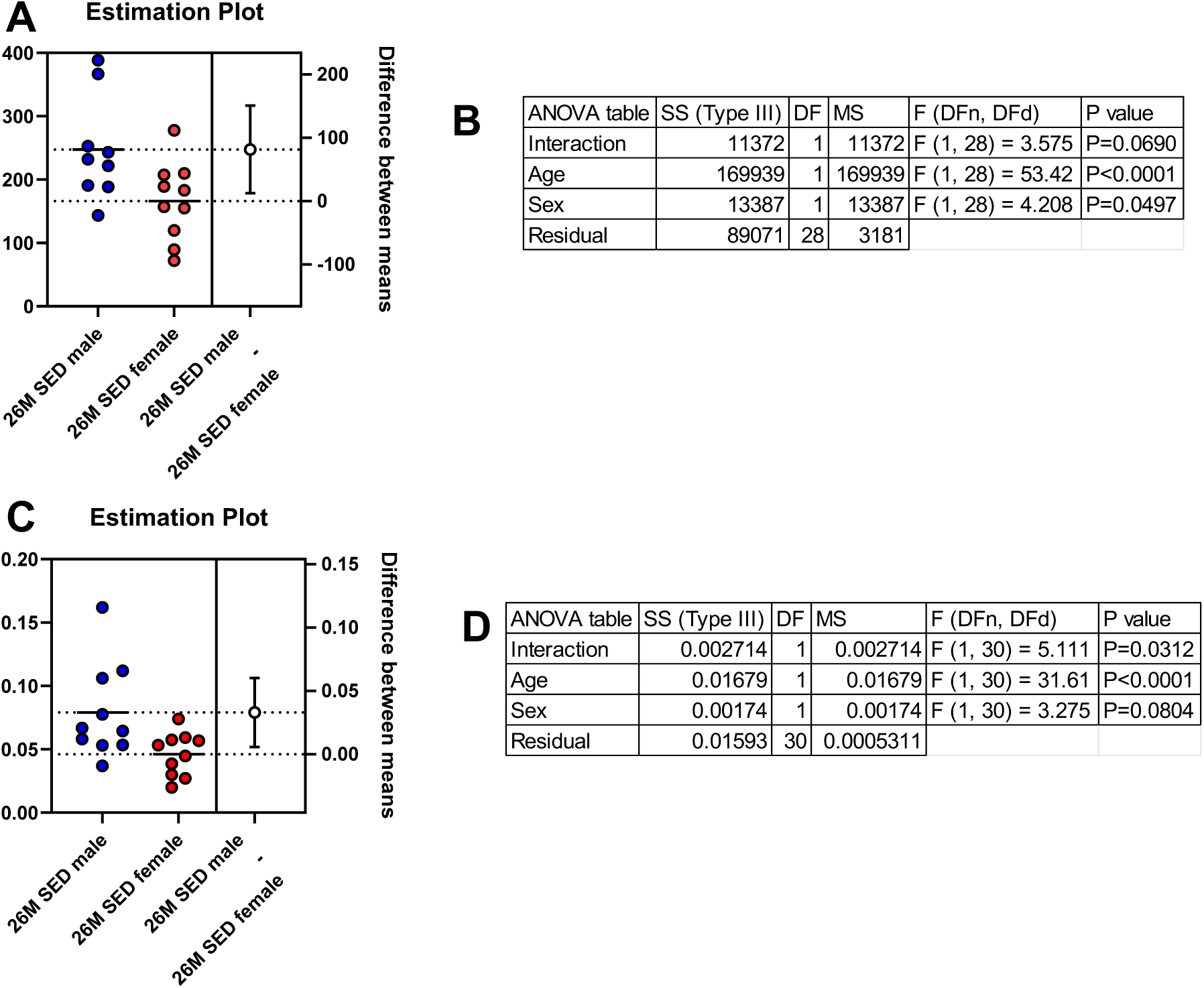
Sex is a significant influence on age-associated increase in mitophagy. (A) Estimation plot with 95% CI showing mean differences in mitolysosome number between male and female sedentary mice at 26 months. (B) Two-way ANOVA table showing group x sex interactions in mitolysosome number. (C) Estimation plot with 95% CI showing mean differences in mitolysosome number normalised to fibre area between male and female sedentary mice at 26 months. (D) Two-way ANOVA table showing group x sex interactions in mitolysosome number normalised to fibre area.

**Extended data figure 4.**
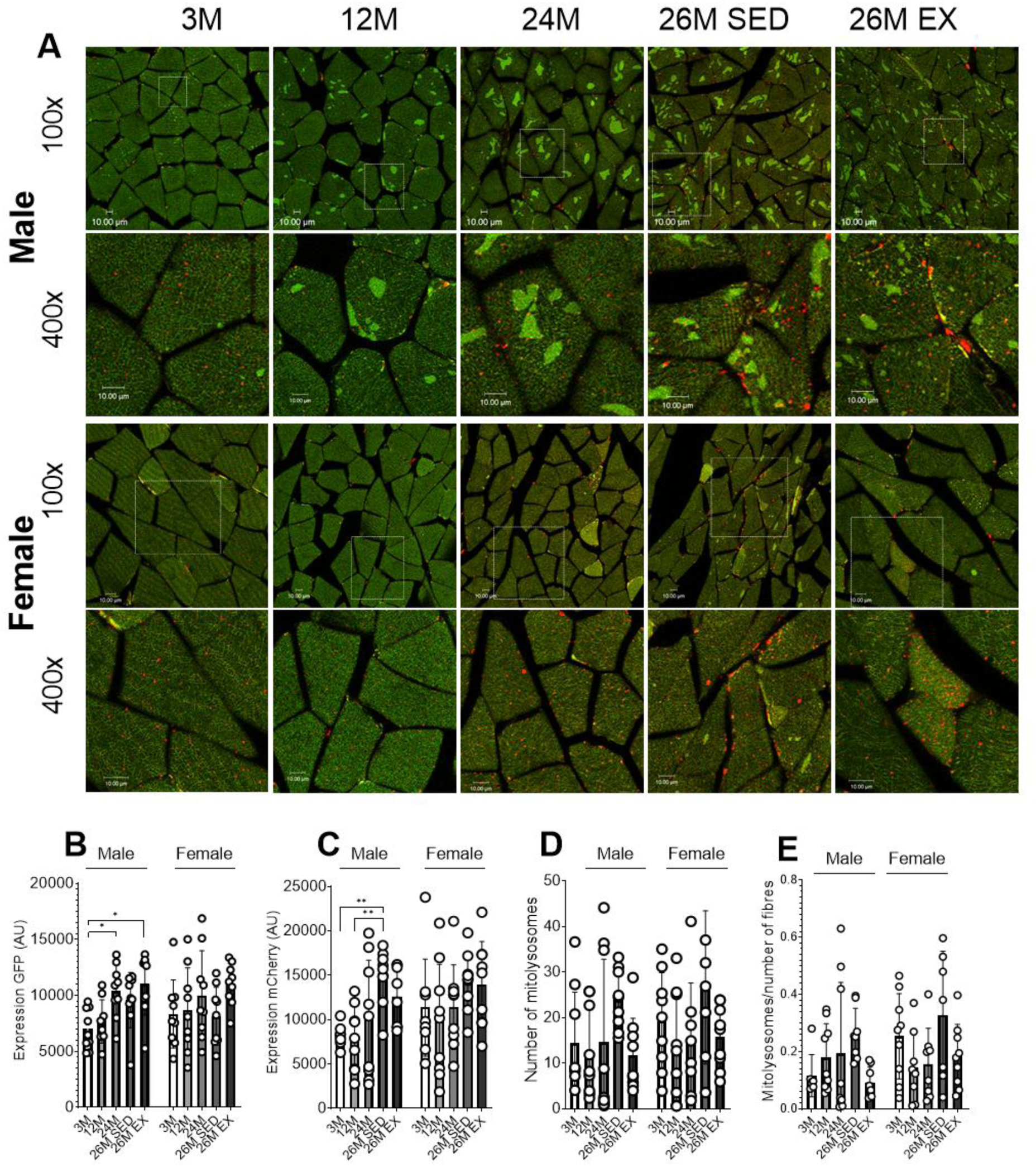
Immunofluorescent imaging of cross-sectional gastrocnemius fibres. (A) Representative confocal images of cross-sectional skeletal muscle fibres in male and female mice at 100x- and 400x- magnification. (B) Measure of GFP fluorescent intensity in fibres. (C) Measure of mCherry fluorescent intensity in fibres. (D) Number of mitolysosomes detected per image. (E) Number of mitolysosomes per fibre area (µm^2^) of image. Data represented as mean ± SD (n=6-10). One-way ANOVA with multiple comparisons, *p<0.05, **p<0.01, ***p<0.001, ****p<0.0001

**Extended data figure 5.**
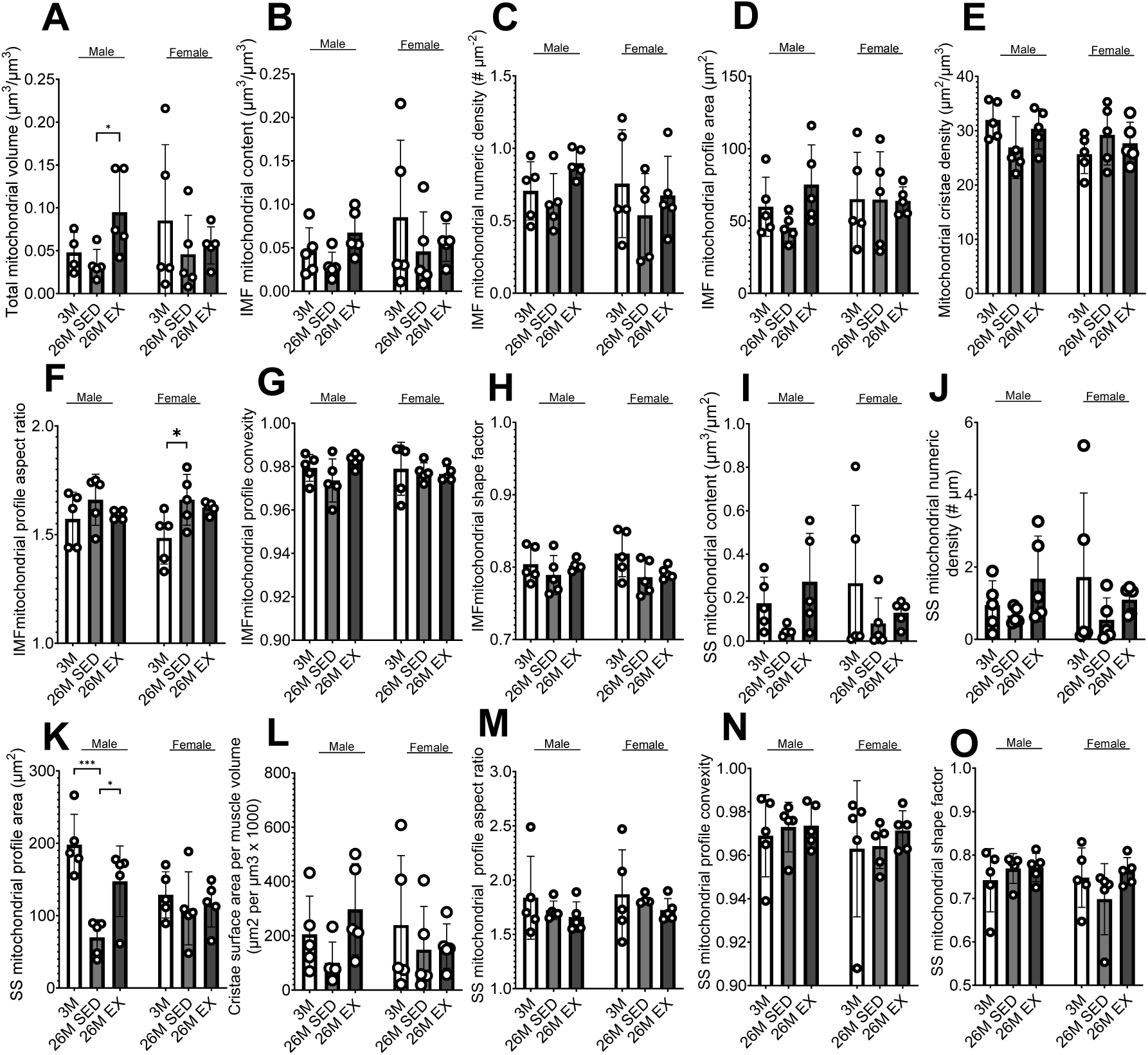
Measure of mitochondrial profiles using transmission electron microscopy. (A) Total mitochondrial volume (µm^3^/µm^3^) in skeletal muscle fibres. (B) IMF mitochondrial content per muscle fibre (µm^3^/µm^3^). (C) Numeric density per image of IMF mitochondria (number mitochondria/µm^2^). (D) IMF mitochondrial profile area (µm^2^). (E) Mitochondrial cristae density. (F) Aspect ratio of IMF mitochondria. (G) Convexity of IMF mitochondria. (H) Shape factor of IMF mitochondria. (I) SS mitochondrial content per fibre surface area (µm^3^/µm^2^). (J) Numeric density per image of SS mitochondria (number mitochondria/length of fibre (µm)). (K) SS mitochondrial profile area (µm^2^). (L) Composite measure of mitochondrial volume density and cristae density providing an estimate of cristae surface area per muscle volume (µm^2^/µm^3^ x 1000). (M) Aspect ratio of SS mitochondria. (N) Convexity of SS mitochondria. (O) Shape factor of SS mitochondria. Data represented mean ± SD (n=5). One-way ANOVA with multiple comparisons, *p<0.05, **p<0.01, ***p<0.001, ****p<0.0001.

